# Septin filament assembly assist the lateral organization of membranes

**DOI:** 10.1101/2024.03.19.585775

**Authors:** Fatima El Alaoui, Isabelle Al-Akiki, Sandy Ibanes, Sébastien Lyonnais, David Sanchez-Fuentes, Rudy Desgarceaux, Chantal Cazevieille, Marie-Pierre Blanchard, Andrea Parmeggiani, Adrian Carretero-Genevrier, Simonetta Piatti, Laura Picas

## Abstract

Compartmentalized interactions of plasma membrane components are essential to support many cell functions, from signaling to cell division, adhesion, migration, or phagocytosis. Cytoskeletal-membrane interactions play an important role in forming membrane compartments, and this feature has been primarily studied through the actin cytoskeleton. Unlike actin, septins directly interact with membranes, acting as scaffolds to recruit proteins to specific cellular locations and as structural diffusion barriers for membrane components. However, how septins interact with and remodel the local membrane environment is unclear. Here we combined minimal reconstituted systems based on fluorescence microscopy and quantitative atomic force microscopy together with live yeast cell imaging and STED microscopy to study septin-mediated membrane organization. Our results show that septins self-assembly into filament-based sub-micrometric patches and high-order structures prompt their membrane-organizing role *in vitro* and in yeast cells, respectively. Furthermore, we show that the polybasic domain of Cdc11, in addition to the amphipathic helix of Cdc12, plays an essential role in supporting the membrane remodeling and curvature-sensing properties of yeast septins. Collectively, our work provides a framework for understanding the molecular mechanisms by which septins can support cellular functions intimately linked to membranes.

## INTRODUCTION

The architecture of the cell cortex is essential to support many cellular processes, such as cell division, trafficking and motility (1, 2). In eukaryotic cells, the cortex is made of a dynamic composite encompassing the plasma membrane and its interacting proteins that is linked to the underlying cytoskeletal network. This cytoskeleton-membrane adhesion has two major roles: mechanical, to support changes in cell shape, for instance during cell blebbing or filopodia protrusion, and biochemical, as it provides a spatial and temporal organizing frame for membrane-associated components (3–5). In the latter case, the cytoskeleton can act as a microscopic structural diffusion barrier for membrane components, as reported for neurons, the base of cilium or the cleavage furrow of dividing cells (5–7). Furthermore, the association of trans-membrane proteins with the underlying actin meshwork is involved in the mesoscale-domain compartmentalization of the cell membrane (5, 8). *In vitro*, the actin cytoskeleton was shown to influence the phase behavior and microdomain formation at membranes depending on the type of anchoring of lipids and proteins (9), the effect of actin polymerization (10), and the stress induced by molecular motors such as myosin (11). The cytoskeleton-mediated compartmentalization of the membrane appears essential to facilitate biomolecular reactions (12, 13) and thus, may be very important for cellular signaling (8, 14). For instance, phosphoinositide signaling, including the receptor G protein/PLC pathway and the PI4K pathway, appears concentrated at liquid-ordered (Lo) domains enriched with cholesterol (14). Also, the activation of type I tyrosine kinase epidermal growth factor (EGF) receptor appears to depend on the cortical actin organization and to polarize at specific cellular locations (13). The type of cytoskeleton cross-linking to the plasma membrane plays an important regulatory role in the formation of specialized membrane compartments (7). For instance, in the case of actin, the interaction with the membrane is not direct but mediated by cytoplasmic proteins, such as the ezrin, radixin and moesin (ERM) family, myosin 1b, anillin and septins (15–20) that bind both actin and lipids, typically phosphatidylinositol-4,5-bisphosphate(PI(4,5)P_2_) (21). In the case of septins, their organization into octamer-based filaments is essential for their role as actin-membrane linkers in human cells (20). Furthermore, the density and adhesion strength of the cytoskeletal linkers are crucial parameters to prevent the cell cortex slippage during mechanically active processes, such as actomyosin contraction, and allow sustained membrane deformations (22–24).

Septins, the fourth cytoskeleton component, might execute a central role in stabilizing the cell cortex architecture during actomyosin contraction. Furthermore, septin organization at the collar of dividing yeast cells was shown to scaffold actomyosin ring assembly for cytokinesis (25–28).

Septins are a family of cytoskeletal proteins highly conserved in animal cells and fungi. They share a common structural organization consisting of a core GTP-binding domain, a septin-unique element and variable N- and C-terminal extensions (29, 30). Septins form heteromeric complexes composed by pairs of different septins and higher-order structures, such as non-polar filaments and rings. These septin structures can be localized at cellular protrusions, such as at the base of cilia and dendritic spines or the bud neck of budding yeast cells, where they act as scaffolds to promote protein-protein interactions and to maintain subcellular compartmentalization by forming diffusion barriers for membrane-associated proteins (6) and, recently, for PI(4,5)P_2_ lipids (31). Septins have been shown to interact with actin (18, 32), microtubules (33, 34) and to associate with actin-binding proteins, such as anillin and non-muscle myosin II (32, 35). Furthermore, septins were shown to bind membranes and this interaction influences, in turn, septin organization into higher-order structures (17, 36). Septins interact with membranes containing anionic lipids, such as PI(4,5)P_2_, phosphatidylserine and cardiolipin (17, 37–39), and display a strong affinity for micron-scale membrane curvatures (40–43). Truncations within the N-terminal domain of septins, which contains a polybasic amino-acid region, showed its role for phosphoinositide selection and membrane association (37, 44–46). In addition, the C-terminal amphipathic helix of Cdc12, which is responsible for septin-curvature sensing (38, 40, 41), was recently reported to facilitate membrane interaction *in vitro* (47).

Because the lipid composition and topology of cellular membranes is heterogeneous, these observations suggest that the appearance of specific septin organizations might be sensitive to the local membrane environment. However, it is unclear if all septin architectures or specific ones would, in turn, affect the lateral organization and compartmentalization of membranes.

We use a combination of time-lapse and super-resolution imaging, atomic force microscopy, and nanotechnology approaches on *in vitro* reconstituted systems and yeast cells to show that membrane-diffusive septins organize into filament-based sub-micrometric patches. These structures, in coexistence with high-order septin structures, directly modulate the lateral organization of membranes. We provide experimental evidence supporting that two membrane-interacting interfaces of septins, the polybasic motif of Cdc11 and amphipathic helix of Cdc12, are involved in this process.

## RESULTS

### Septins regulate the lateral organization of the budding yeast membrane during the cell cycle

In many eukaryotic cells, one of the central and most studied functions of septins is to promote cytokinesis, i.e. the last step of cell division (48). In budding yeast, the cytokinetic function of septins is linked to their unique ring-like organization at the bud neck and their interaction with the plasma membrane (28). To investigate the role of septins in membrane organization at the bud neck of *S. cerevisiae*, we monitored the general polarization (GP) of the lipophilic probe Laurdan as a readout of the lipid packing of the plasma membrane (Figure 1). Briefly, the GP quantitatively reports the shift in the emission spectrum of the probe depending on the polarity of its immediate lipid environment. Thus, an increase in the environment polarity is typically associated with liquid-disordered (Ld) phases, being more fluid and less packed, as compared to liquid-ordered (Lo) phases. The relative GP index, between −1 and 1, is inversely proportional to the membrane polarity and consequently, it numerically increases with lipid packing or order (49, 50). We used yeast cells expressing Cdc3 endogenously tagged with mCherry at its N-terminus (mCherry-Cdc3) to correlate GP values with septin localization in early G1, when the septin signal is absent from the presumptive bud site/bud neck, and during the rest of the cell cycle, where septins are bound to the plasma membrane at the bud neck and form a collar (Figure 1B). We observed a heterogenous lateral organization of GP values at the plasma membrane of yeast cells, as denoted by the presence of patches with high GP values (GP > 0), irrespective of the cell cycle phase (Figure 1B). The segmentation of the cell cortex from transmitted light images indicated, however, that the overall GP values at the yeast plasma membrane was relatively constant throughout the entire cell cycle (Figure 1C). The angular cross-section analysis (Figure 1D) of the mCherry-Cdc3 signal and GP values at the cortex of mother cells showed that the presence of septins at the bud neck correlates with a local increase in the Laurdan GP index (∼DGP ∼ 0.3-0.5), as compared to septin-lacking regions either at early G1 (unbudded cells lacking a septin ring) or during the rest of the cell cycle (budded cells) (Figure 1E). We observed that while the septin signal spanned a region of ∼1 μm, in agreement with the expected bud neck diameter (26), reorganization of GP values expanded over larger distances, suggesting that septins might affect the lateral membrane fluidity over a long-range. To asses that the effect was not due to actin cables and patches that also populate the bud neck (51), we treated cells with Latrunculin A (LatA), which inhibits actin polymerization (Figure 1C-E). Under these conditions, we observed the same trend in the local increase of GP values overlapping with the septin signal at the division site without affecting the global fluidity of plasma membrane, suggesting that the observed effects are independent of the actin cytoskeleton organization and likely ascribable directly to septins.

**Figure 1.**
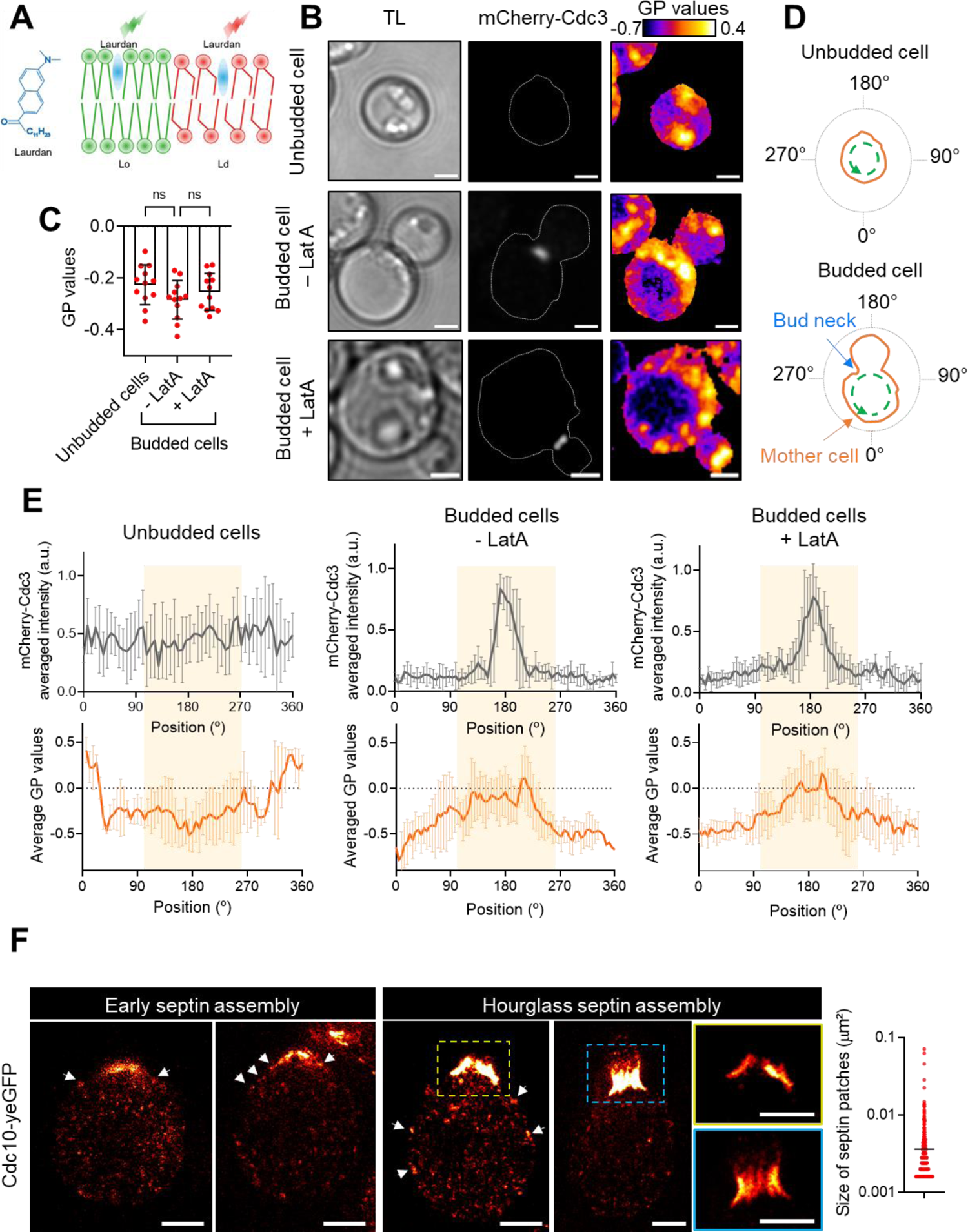
Septin architecture and phase behavior of the yeast plasma membrane during the cell cycle. **A)** Schematic representation of the lipophilic probe Laurdan (blue) and its partitioning in Lo (green) and Ld phases (red). **B)** Transmitted light (TL) and confocal images of yeast cells expressing mCherry-Cdc3 (gray) and stained with the Laurdan probe after 10 min incubation at 30°C. Color-coded GP images (fire LUT) of z-stacks max-projections of unbudded or budded cells, either untreated (DMSO, -LatA) or treated with 0.1 mM of LatrunculinA (+LatA). Scale bar, 2 µm. **C)** Quantification of the global GP index at the plasma membrane of unbudded or budded cells, either untreated (DMSO, -LatA) or treated with 0.1 mM of LatrunculinA (+LatA). Number of cells analyzed, n = 12, 12 and 12, respectively, from at three biological replicates. Error bars represent s.d.; Mann-Whitney test: n.s. P > 0.05. **D)** Schematics showing the angular cross-section analysis (green-dashed line) of the plasma membrane (orange) starting for mother cells at the opposite pole of the bud neck or, for unbudded cells, at an arbitrary position, 0°, following a counterclockwise direction. **E)** Average cross-section analysis (n = 6 cells) showing the mCherry-Cdc3 signal (gray) and GP index values (orange) at different angular positions, as detailed in D from three biological replicates. Error bars represent s.d. **F)** STED images of yeast cells expressing Cdc10-yeGFP revealed with an anti-GFP nanobody Atto647N during early septin assembly at the presumptive bud site and early hourglass assembly at the bud neck. Scale bar, 1 µm. Distribution of the apparent area of septin patches (μm^2^) at the mother cell. N = 10 cells, number of patches, n = 421 from two biological replicates.

Septin architecture and membrane remodeling are intimately connected during the cell cycle (28). Consequently, we asked if specific septin architectures may correlate with the observed effect on the lateral organization of the yeast plasma membrane. To this end, we performed STED microscopy on yeast cells endogenously expressing Cdc10-yeGFP and immunolabeled with an anti-GFP nanobody Atto 647N (52). Under our spatial-resolution (i.e., ∼ 45 nm), we could detect an accumulation of the Cdc10-yeGFP signal at the presumptive bud site at early stages of septin assembly that turned into a filament-like organization upon septin-hourglass formation (Figure 1F), in line with a previous report (52). Interestingly, we also identified the presence of discrete sub-micrometric septin patches that were particularly visible at the expected contour of the cell and, often, in proximity to high-order septin structures (Figure 1F and Figure 1-figure supplement 1), which shows the negative control, i.e. yeast cells not expressing Cdc10-yeGFP). These observations suggest that high-order and sub-micrometric septin structures at the plasma membrane might potentially contribute to septin-mediated lateral membrane organization.

### Assembly of septin patches on membranes is a diffusion-driven process

To get insights into the mechanisms leading to septin patch formation in yeast cells, we set out to investigate *in vitro* the contribution of the lipid membrane, as this feature was previously shown to modulate septin architecture (17, 37). First, we determined by liposome floatation assays the binding affinity of septins for membrane lipids at physiological ionic strength (i.e. 150 mM NaCl). To this end, we used purified recombinant yeast septin octamers (Cdc11-Cdc12-Cdc3-Cdc10-Cdc10-Cdc3-Cdc12-Cdc11), as previously reported (53), and we allowed their binding to small unilamellar vesicles (SUVs) made of either 100% mol phosphatidylcholine, as a neutral lipid mixture, or 25% of total negatively charged lipids, including phosphatidylinositol (PI) or equivalent % mol of PI(4,5)P_2_, phosphatidic acid (PA), or phosphatidylserine (PS) at the expenses of PI, as detailed in Table S1. We determined septin binding through western-blot analysis by comparing the signal of Cdc11 at the top (SUV-bound protein) and bottom (unbound protein) of a sucrose gradient (54) (Figure 2A). In the absence of SUVs, septins were collected in the bottom fraction (CTRL, Figure 2A). In the presence of SUVs, yeast septins bound to SUVs of almost all tested negatively charged lipid compositions, except for PS, with an increased affinity for PI(4,5)P_2_, in agreement with previous studies (17, 37–39). Noteworthy, septins also bound the neutral PC lipid, pointing out that they not only display an affinity for anionic lipids.

**Figure 2.**
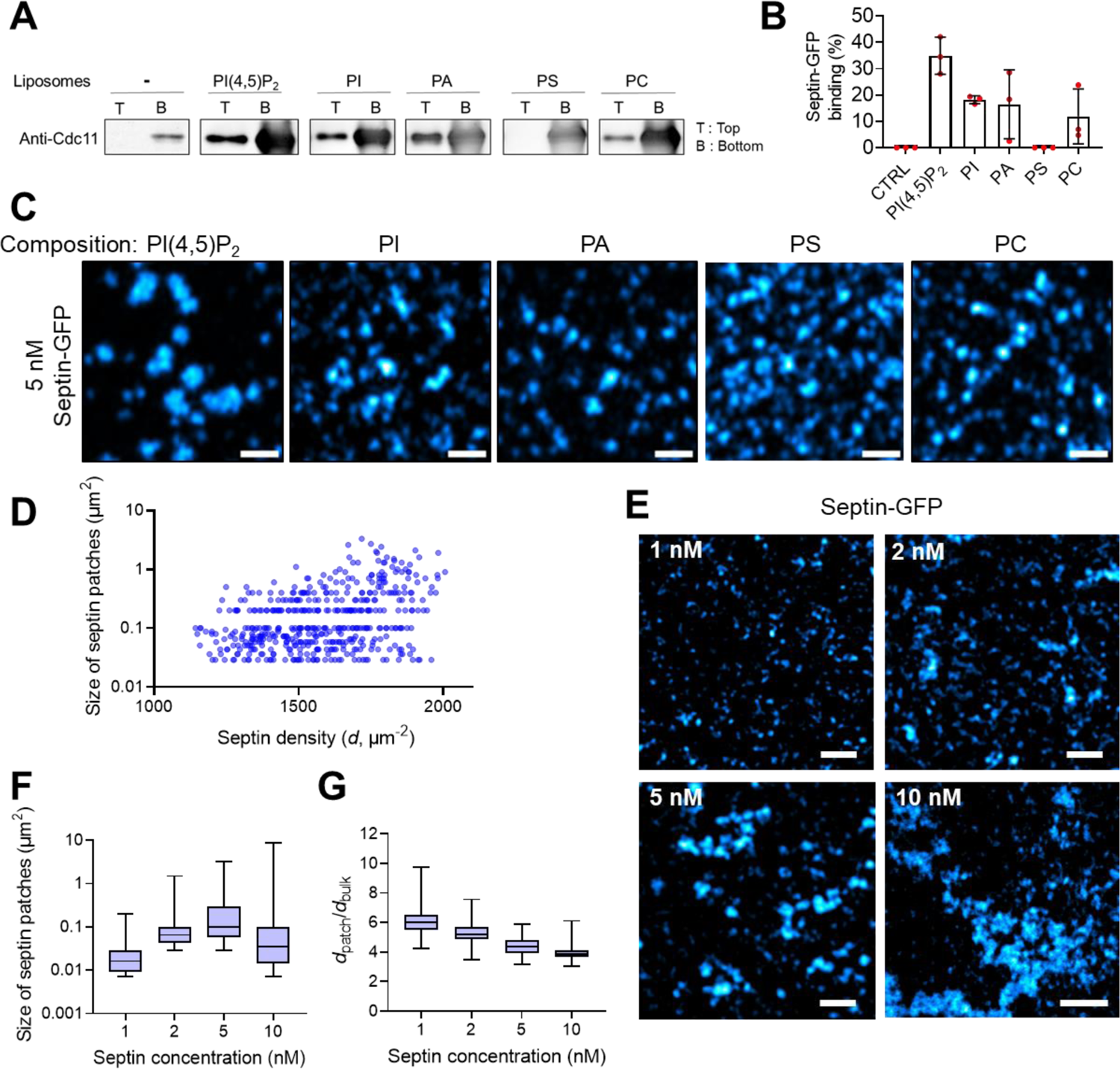
Binding and organization of septins on lipid membranes. **A)** 1 µM of Septin-GFP hetero-octamers were incubated without liposomes (-, CTRL) and with liposomes (2 mM) made of egg-phosphatidylcholine (PC) or with different lipid mixtures containing 25% total negative charge: Liver-phosphatidylinositol (PI), Brain-PI(4,5)P_2_ (PI(4,5)P_2_), Egg-phosphatidic acid (PA), Brain-phosphatidylserine (PS). The suspension was subjected to liposome floatation assays. The composition of each liposome suspension was: PI(4,5)P_2_: 80% mol PC, 15% mol PI, 5% mol PI(4,5)P_2_; PI: 75% mol PC, 25% mol PI; PA: 75% mol PC, 15% mol PI, 10% mol PA; PS: 75% mol PC, 15% mol PI, 10% mol PS; PC: 100% mol PC. 1 µM of Septin-GFP hetero-octamers were polymerized in septin polymerization buffer and the liposome-bound (Top, T) or free (Bottom, B) septin Cdc11 was detected by western blotting with anti-Cdc11 antibody. **B)** Quantification of septin-GFP binding from liposome floatation assays (right panel): septin-GFP binding = Top/(Top+Bottom) x 100. Results represent the mean ± s.d. from three biological replicates. **C)** Airyscan images showing the association of 5 nM septin-GFP in septin buffer with SLBs made with the same lipid composition as in A. Scale bar, 1 μm. **D)** Representation of the area of septin patches (μm^2^) as a function of the septin density (*d*, μm-^2^) on 2.5% mol PI(4,5)P_2_-SLBs upon addition of 5 nM of septin-GFP heterocomplexes (n = 477 septin patches, from three biological replicates). **E)** Airyscan images showing the association of 1 nM, 2 nM, 5 nM and 10 nM of septin-GFP with SLBs made of 2.5% mol PI(4,5)P_2_. Scale bar, 2 μm. **F)** Representation of the area of septin patches (μm^2^) as a function of septin concentration (nM). **G)** Ratio of the membrane-bound septin density (*d*, μm-^2^) on patches (*d*_patch_) versus at the membrane bulk (*d*_bulk_) as a function of septin concentration (nM). Results in F and G represent the mean ± s.d (n = number of patches from two biological replicates, n = 615, 351, 477, 1036, respectively).

We then characterized by sub-diffraction Airyscan microscopy the contribution of the lipid composition to the spatial organization of septins (Figure 2C). To this end, 5 nM of GFP-tagged septin octamers were allowed to bind to supported lipid bilayers (SLBs) made with the same composition as that used in liposome floatation assays, with the addition of trace amounts (0.2%) of Atto647N-labeled DOPE (A647-DOPE) to visualize the bilayer surface (Figure 2-figure supplement 1). Airyscan imaging showed that septins organized into sub-micrometric patches randomly distributed over the bilayer surface for all tested lipid compositions (Figure 2C). The formation of septin patches was, however, more evident on PI(4,5)P_2_-containing SLBs.

To get further insights into the formation of septin patches, we focused on SLBs containing 2.5% mol PI(4,5)P_2_, as it was the lipid composition displaying the highest septin association with membranes (Figure 2A). Furthermore, PI(4,5)P_2_ is predominantly localized at the plasma membrane, where many of the septin’s functions have been reported to take place (29, 30). We estimated the apparent area of septin patches on PI(4,5)P_2-_containing SLBs to be between 0.05 to 2 μm^2^ (Figure 2D), in agreement with the same range of magnitudes observed in yeast cells (Figure 1F). The quantification of the density of membrane-bound GFP-tagged septin octamers (*d*), as described in (55, 56), showed an increased density of septins in larger patches (Figure 2D). However, we also detected a substantial population of sub-micrometric septin patches displaying high septin densities. This observation suggests that septin patches might contain layers of septin octamers, typically occurring in a dense septin meshwork, in agreement with a recent report (39).

Next, we evaluated the contribution of septin concentration by monitoring septin patch formation upon addition of 1 nM, 2 nM, 5 nM, or 10 nM of GFP-tagged septin octamers on SLBs made with 2.5% mol PI(4,5)P_2_ (Figure 2F-G). We observed that the apparent area of septin patches increased with septin concentration up to 10 nM, where septin patches started to connect into a large and single structure. Evaluation of the density of septin octamers forming patches relative to the density of bulk membrane-bound septins (*d*_patch_/*d*_bulk_) showed a monotonic decrease with increasing septin concentrations (Figure 2G). Therefore, membrane-bound septin octamers preferentially assemble into patches rather than diffusing in the membrane bulk at low septin concentration.

The formation of elongated septin filaments and high-order structures has been proposed to stem from a diffusion-driven motion of septins in the presence of membranes (57). The results above indicate that septin patch formation might also originate from a 2D diffusion of septin octamers on membranes. To get further insights into the contribution of this process, we monitored the fluorescence recovery of GFP-tagged septin signal on PI(4,5)P_2_-containing SLBs upon the addition of 5 nM of septins and formation of septin patches (Figure 3A-C). Upon septin patch bleaching, as indicated by the cross-section analysis in Figure 3B (orange dashed line in Figure 3A), we observed a mild fluorescence recovery of the septins-GFP signal after ≥ 7 min on septin patches (Figure 3C). Indeed, we obtained that the recovery time of the fluorescent signal, τ, was τ = (77 ± 12)*s*. We can then estimate the mobile fraction 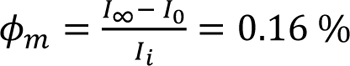 and the immobile fraction Φ_*imm*_ = 1 – Φ_*m*_ = 0.84 %. High immobile fraction indicates a high effective viscosity in the septin patch. An effective diffusion constant can be estimated by *D_eff_ = ω^2^/4τ ∽ (0.99 ± 0.16)10^-14m2^/S* indicating a slow two-dimensional septin dynamics (Figure 3-figure supplement 1). This value was still more than a decade lower than the isotropic diffusion constant (*D*_i_) of a single septin-octamer center of mass at long time (58):

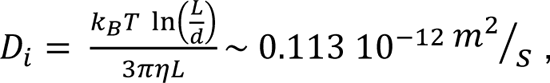

where *L* ∼ 32 nm is the septin octamer length, *d* ∼ 4 nm is the rod diameter, *η* ∼ 1 Pa.s is the membrane viscosity (1000 times the water viscosity), *k*_*B*_ = 1.38 10^−23^*J/K* is the Boltzmann constant and *T* ∼ 300 *K* is the system temperature (in Kelvin units). We confirmed that this trend was independent of the SLB fluidity by evaluating the fluorescence recovery of the lipid membrane (Figure 3-figure supplement 2). The cross-section analysis of the septin signal at ≥ 7 min post-bleaching showed a moderate increase in the septin intensity at the bleached region relative to the same region at the moment of bleaching (Figure 3D), suggesting that the diffusion of septin octamers in the membrane bulk may contribute to the formation of the septin patch. To confirm the involvement of a diffusion-driven process in septin-patch assembly, we depleted the system from membrane-unbound septins by washing out three times the imaging chamber. Under these conditions, we could not detect a decrease in the membrane-bound septin-GFP signal both in the bilayer bulk and septin patch, suggesting that depletion of septins in solution do not affect the behavior of septin patches once the equilibrium is reached (Figure 3-figure supplement 3). As expected, replenishment of the system with 5 nM of GFP-tagged septin complexes was accompanied by a rise in the global septin intensity that led, after 7 min of incubation, to a significant increase of the intensity at the bleached and non-bleached region of the septin patch. This observation suggests that the assembly of septin octamers into a patch-like structure might be driven by septin diffusion in the membrane plane and the addition of septin layers, possibly through the association of septins from the bulk solution.

**Figure 3.**
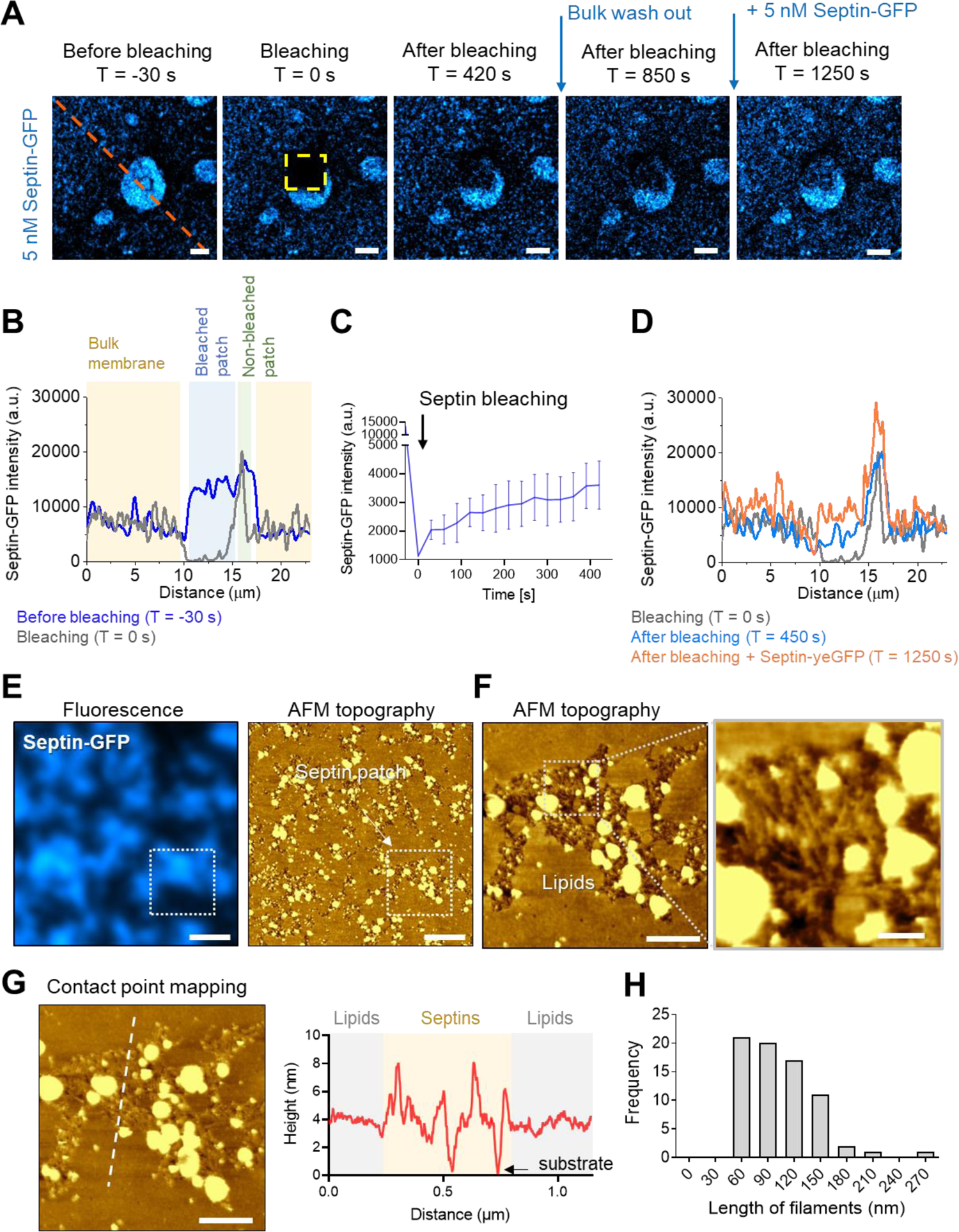
Dynamics and structural organization of septins on PI(4,5)P_2_-containing SLBs. **A)** Confocal images of septin-GFP bleaching and fluorescence recovery experiment after 10 min binding of 5 nM of septin-GFP on 2.5% mol PI(4,5)P_2_-containing SLBs at different time points: before bleaching (T = −30 s), at the bleaching moment (T = 0 s), after bleaching (T = 420 s), after the solution wash out (T = 420 s), and after replenishment with 5 nM of septin-GFP (T = 1250 s). The FRAP ROI is indicated in yellow. Scale bar, 2 μm. **B)** Cross-section of the septin-GFP signal along the orange line in A before bleaching (blue) and at the bleaching moment (gray). The different features of the septin signal on SLBs are highlighted (septins at the bulk membrane and bleached and non-bleached septin patch). **C)** Representation of the septin-GFP intensity as a function of time (in seconds) at the FRAP region. Graph represents the mean ± s.d (n = 8, ROIs from two independent experiments). **D)** Cross-section of the septin-GFP signal along the orange line in A at the bleaching moment (gray), after bleaching (blue), and after replenishment with 5 nM of septin-GFP (orange). **E)** Correlative fluorescence (left) and AFM topography (right) images of septin-GFP patches on 2.5% mol PI(4,5)P_2_-SLBs. Scale bar, 1 µm. False-color scale in AFM images, 17 nm. **F)** Magnified AFM topography images corresponding to the white box in E (scale bar, 250 nm and 50 nm for the magnified image. False-color scale, 7 nm and 20 nm). **G)** Contact point image of the topography image in F (scale bar, 250 nm. False-color scale, 7 nm). Cross-section along the white line showing the height of septin-GFP hetero-octamers and the membrane relative to the glass substrate. **H)** Length distribution of septin filaments (n = 73 from two independent AFM experiments).

To get further insight on the structural organization of septins on PI(4,5)P_2_-containing SLBs, we used correlative light and atomic force microscopy (CLAFM) (Figure 3E-G). Addition of 5 nM of GFP-tagged septin octamers to SLBs lead to the formation, as expected, of septin patches that we first identified using fluorescence microscopy and then characterized through AFM topography imaging (Figure 3E). In agreement with previous AFM studies, we also observed the appearance of lipid blobs associated with yeast septins (47, 53), possibly resulting from bacterial lipids co-purifying with septins, as previously reported for other lipid-binding proteins (59). The cross-section analysis of the contact point map, which corresponds to the sample’s topography acquired at the minimal loading force (i.e., at the tip-sample contact point), showed that septin patches mostly protrude by ∼ 4 nm above the lipids (Figure 3F). This height is consistent with the thickness of yeast septin octamers (60), suggesting that septins are likely laying as a monolayer on top of the membrane. Furthermore, the presence of some peaks with an 8 nm height suggests that septins can also be present as two layers on top of each other, in agreement with previous studies with *Drosophila* septins (39). Closer inspection of septin patches showed that they are formed of randomly organized short septin filaments of a length between 60 to 150 nm (Figure 3G), which corresponds to 2 to 5 multiples of octamers (60). Collectively, these observations suggest that yeast septin hetero-octamers assemble as filament-based patches on PI(4,5)P_2_-containing membranes.

### Septin assemblies modulate the microscopic organization of membranes *in vitro*

Next, we set out to investigate if filament-based septin assemblies observed on PI(4,5)P_2_ membranes might also impact lipid organization *in vitro*. To this end, we monitored the binding of 5 nM GFP-tagged yeast septin octamers onto 2.5% mol PI(4,5)P_2_-containing SLBs doped with 0.2% mol of two distinct lipid dyes: Atto647N-labeled DOPE (A647-DOPE) and Texas Red-labeled PI(4,5)P_2_ (TF-TMR-PI(4,5)P_2_) (Figure 4A-B). Sub-diffraction Airyscan microscopy images showed that before septins addition, SLBs displayed a homogenous distribution of both lipid dyes. As expected, addition of 5 nM of septins led to the immediate formation of septin domains that was accompanied by a marked time-dependent redistribution of the A647-DOPE dye (Figure 4A and Figure 4-figure supplement 1). Cross-section analysis showed that A647-DOPE-depleted regions co-localize with the occurrence of septin patches (Figure 4B). Conversely, at the same regions, we did not detect an apparent depletion of the TF-TMR-PI(4,5)P_2_ dye. The kymograph analysis of septin binding also showed that nucleation and growth of septin patches immediately engage a selective depletion of the DOPE dye with respect to the PI(4,5)P_2_ dye (Figure 4C). The analysis of the relationship between the apparent size of septin patches as a function of the average intensity of either the A647-DOPE or TF-TMR-PI(4,5)P_2_ signal showed that the intensity of the PI(4,5)P_2_ dye appeared steady and thus, independent of the septin patch size (Figure 4D). Conversely, depletion of the DOPE signal showed a size-dependency, with an almost total signal depletion with septin patches larger than 0.1 µm^2^. We found that depletion of the A647-DOPE signal induced by the septin patches was not due to fluorescence quenching (Figure 4-figure supplement 2), as we did not detect an increase of the Atto647N signal upon bleaching GFP-tagged yeast septins. Furthermore, A647-DOPE depletion did not depend on the wavelength and type of the fluorescent dye of the DOPE derivative (Figure 4-figure supplement 3). We also confirmed that the effect on A647-DOPE partitioning was specific to septins by monitoring the lipid organization upon addition of 1 μM of the PH domain of PLCδ1 (PH^PLCδ1^), which selectively interacts with PI(4,5)P_2_ (61), fused to GFP (GFP-PH^PLCδ1^) on 2.5% mol PI(4,5)P_2_-containing SLBs (Figure 4-figure supplement 4). We found that addition of GFP-PH^PLCδ1^ led to its immediate membrane association. However, this was not mirrored by any significant lateral redistribution of the A647-DOPE or TF-TMR-PI(4,5)P_2_ lipid dyes (Figure 4-figure supplement 4). Collectively, these data suggest a direct effect of septins on the lateral distribution of lipids. This feature appears to be selective for the A647-DOPE reporter and moderate or undetectable for other lipid derivatives, such as PI(4,5)P_2_ or phosphatidylcholine (PC) (Figure 4-figure supplement 3C-D). This septin-mediated selective exclusion of A647-DOPE molecules in the membrane plane might be related to the decreased affinity of septins for phosphatidylethanolamine (PE) observed in liposome floatation assays at different % mol of PE (Figure 4-figure supplement 5).

**Figure 4.**
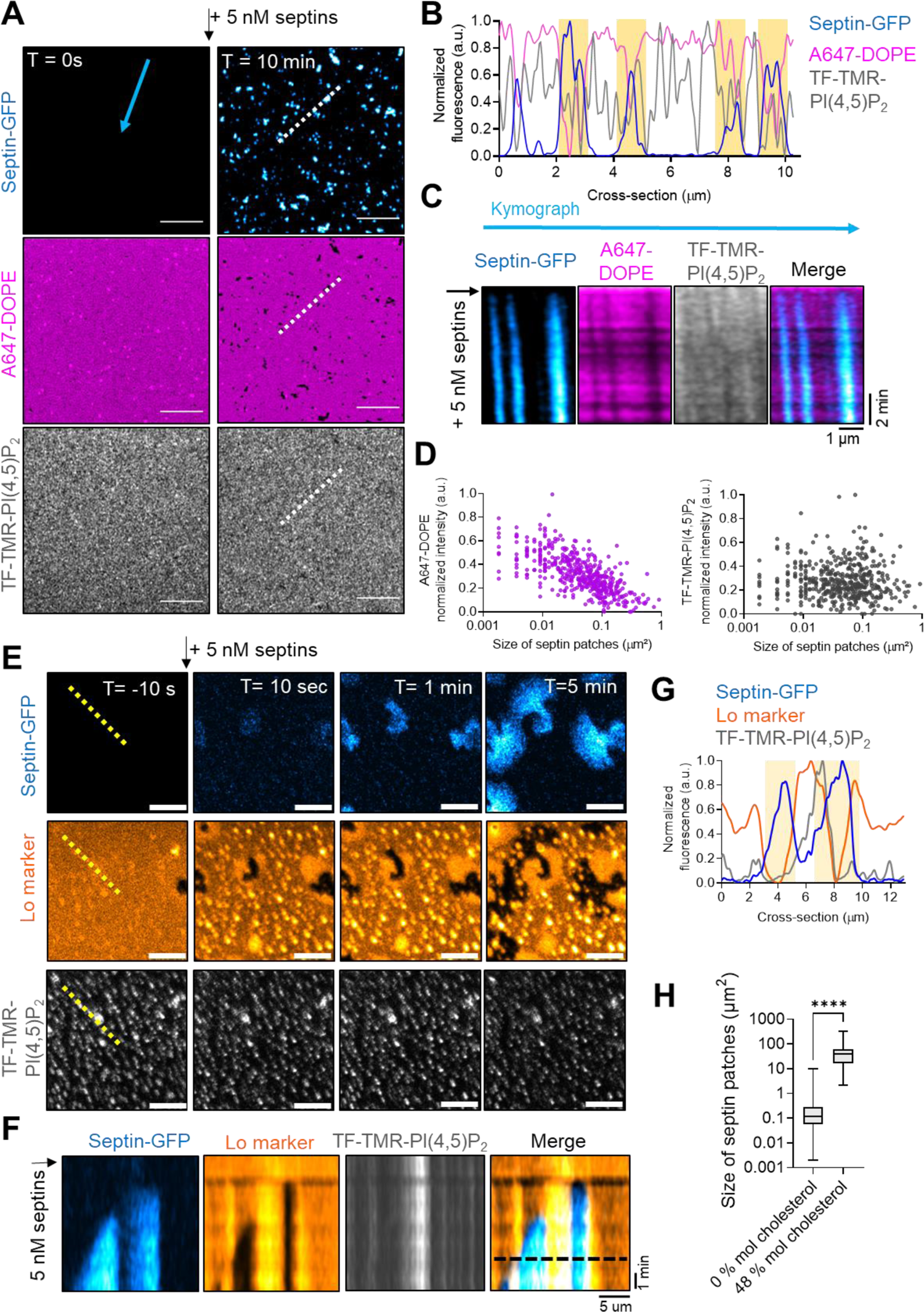
Microscopic organization of *in vitro* membranes upon septin binding. **A)** Airyscan images showing the organization of 5 nM septin-GFP hetero-octamers (in blue) injected in septin buffer onto SLBs containing 2.5% mol Brain-PI(4,5)P_2_ and doped with 0.2% mol fluorescent TF-TMR-PI(4,5)P_2_ (in gray) or 0.2% mol fluorescent A647-DOPE (in magenta) at T = 0 and T = 10 min. Scale bar, 5 μm. **B)** Cross-section analysis along the dashed line in A. **C)** Kymograph analysis along the blue arrow in A of septin-GFP signal (blue), A647-DOPE (magenta) and TF-TMR-PI(4,5)P_2_ (gray) from T = 0 to 8 min. **D**) Intensity distributions of A647-DOPE (magenta) and TF-TMR-PI(4,5)P_2_ (gray) relative to the size of septin patches (μm^2^) from three biological replicates. **E)** Airyscan images showing the dynamics of septin-GFP hetero-octamers (in blue) binding to mica-supported SLBs made of 2.5% mol Brain-PI(4,5)P_2_, 2,5% mol DOPC, 30% mol DPPC, 44.8% mol cholesterol, 20% mol Liver-PI, Lo marker (0.2% mol fluorescent DSPE-PEG-ASR, in orange) and TF-TMR-PI(4,5)P_2_ (in gray). Scale bar, 5 μm. **F)** Kymograph analysis along the yellow line box in E of the septin-GFP (blue), DSPE-PEG-ASR (Lo marker, orange) and TF-TMR-PI(4,5)P_2_ (gray) signal from T= 0 min to 5 min. **G)** Cross-section analysis along the dashed line in the kymograph. **H)** Quantification of the size of septin domains on membranes containing 0% mol or 48% mol cholesterol from Airyscan images. Domains analyzed, n = 6693 and 163, respectively from ≥ three replicates. Error bars represent s.d.; Mann-Whitney test: **** P < 0.0001.

Next, we assessed if septins can directly modify the macroscopic organization of liquid-ordered (Lo) and liquid-disordered (Ld) phases using a model ternary mixture consisting of unsaturated DOPC, saturated DPPC and cholesterol that is typically used to mimic the plasma membrane of eukaryotic cells (9, 49). We monitored the dynamic interaction of 5 nM septin-GFP complexes on mica-supported lipid membranes made of 2.5% mol Brain-PI(4,5)P_2_, 2.5% mol DOPC, 30% mol DPPC, 44.8% mol cholesterol, 20% mol Liver-PI and trace amounts (0.2% mol) of the Lo marker DSPE-PEG-ASR (49) (Figure 4E-G). After mica-SLB preparation, samples were equilibrated at room temperature for one hour and placed under the confocal/Airyscan microscope at a temperature of ∼ 25 °C. Under these conditions, the ternary mixture containing 2.5% mol PI(4,5)P_2_ displayed a mixed phase, as the Lo marker did not appear to display preferential partitioning (Figure 4E). The addition of septin complexes led to their immediate association with membranes and the formation of septin patches with a larger size as compared to SLBs lacking cholesterol (Figure 4H). We observed that cholesterol-containing SLBs septin domains expanded over time and eventually collided, reaching micrometric sizes after ≥ 5 min. The appearance of septin patches was mirrored by a marked reorganization of the Lo marker that appeared excluded from the septin-GFP signal (Figure 4G), suggesting a direct effect of septins on the behavior of Lo/Ld phases *in vitro*. Collectively, these observations point out that formation of septin patches directly modulate the phase behavior and macroscopic re-organization of lipids *in vitro*.

### The polybasic domain of Cdc11 and amphipathic helix of Cdc12 support the membrane reorganization and curvature-sensing properties of septins

Septin interaction with membranes is thought to be mediated by a polybasic (PB) motif, adjacent to the GTP-binding motif at the N-terminus, and by an amphipathic helix (AH) downstream to the septin-unique element (SUE) (17, 37, 40–42, 47, 62). While the PB motif has been primarily reported to mediate septin interaction with anionic lipids (17, 37, 38, 45), the AH was shown to mediate both membrane association and micron-scale curvature sensing of septins *in vitro* and *in cellulo* (40, 41, 43). To determine the contribution of the PB and AH domains on the lipid reorganization-mediated by septins, we generated and purified recombinant GFP-tagged septin octamers carrying a deletion of the PB motif of Cdc11 (residues 2-18; ΔPB-Cdc11) or a C-terminal deletion of Cdc12 that eliminates part of the AH (residues 390-407; ΔAH-Cdc12), or the complete deletion of the C-terminal extension (CTE) of Cdc12 (residues 339-407; ΔCTE-Cdc12) (Figure 5A-E and Figure 5-figure supplement 1). These mutants were designed on the basis of previously published septin mutants (17, 41, 47). First, we assessed by transmission electron microscopy (TEM) after negative staining the ability of the septin mutant proteins to polymerize on 2.5% mol PI(4,5)P_2_-containing lipid monolayers (Figure 5A). We observed that under our conditions wild-type septin-GFP complexes annealed end-to-end, forming short single filaments made of two to three multiples of octamers, as well as long paired filaments connected by regular lateral bridges. Under these conditions, all complexes containing ΔPB-Cdc11 or ΔAH-Cdc12 or ΔCTE-Cdc12 mutants kept their ability to associate with lipid monolayers, although they did not organize into long paired filaments. For the ΔPB-Cdc11 mutant this observation is consistent with the known role of the α0 helix of Cdc11, where the PB motif is located to mediate septin polymerization in solution (60). However, while the Cdc12 mutants assembled into short filaments mostly made of four septin hetero-octamers, the ΔPB-Cdc11 complexes stayed mainly as octamers or double octamers (Figure 5C).

**Figure 5.**
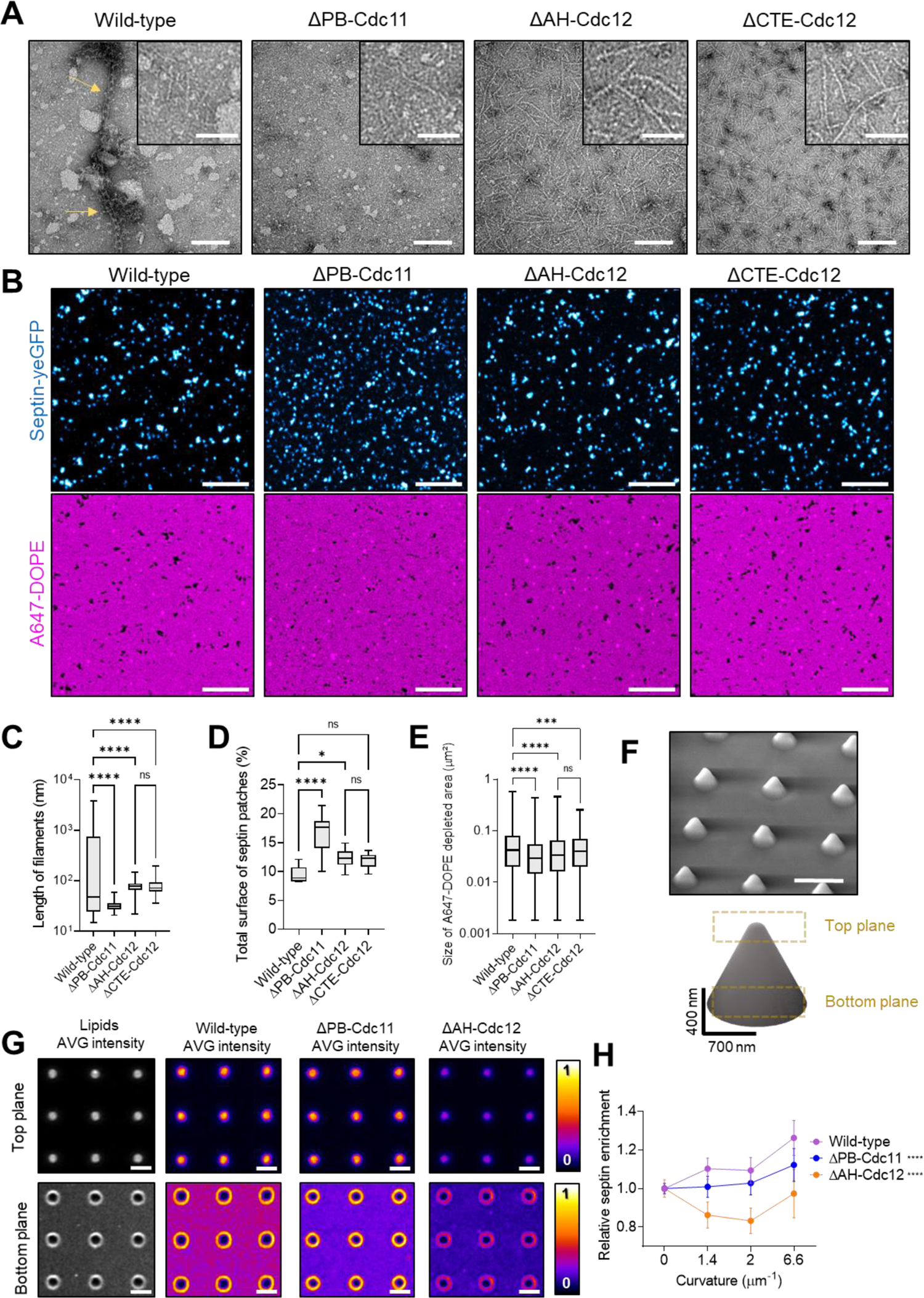
Association of wild-type septins and septin mutants with lipid membranes. **A)** TEM images after negative staining showing the structural organization of 100 nM wild-type septin complexes or ΔPB-Cdc11, ΔAH-Cdc12, ΔCTE-Cdc12 mutant septin complexes on monolayers made of 77.5% mol Egg-PC, 2.5% mol Brain-PI(4,5)P2, 20% mol Liver-PI. Samples were incubated overnight at 4°C. Scale bar, 100 nm, 50 nm for the inset. **B)** Airyscan images showing the association of 5 nM wild-type or mutant GFP-septin hetero-octamers with SLBs containing 2.5% mol PI(4,5)P_2_ and 0.2% mol fluorescent A647-DOPE at T = 1h. Scale bar, 1 μm. **C)** filament length (in nm) of wild-type septins and septin mutants obtained from the TEM images in A (number of filaments, n = 48, n = 33, n = 35, n =50, as in the graph from two independent experiments). **D)** Total surface (%) of the SLB occupied by septin patches of wild-type or mutant septin-GFP hetero-octamers (n = 10 images analyzed per condition, from three biological replicates). **E)** Size of A647-DOPE-depleted regions after addition of wild-type and mutant septin-GFP hetero-octamers (n = 4790, 5464, 450, 5425, respectively (as in the graph order) from three biological replicates). **F)** SEM images of silica substrates displaying an array of cones with a radius r ∼ 0.7 µm at their base and a height h ∼ 800 nm generated by soft-NIL. Scale bar, 3 µm. Representation of the acquisition of a single plane at the top and bottom of cones to visualize the distribution of lipids and septins. **G)** Average intensity (AVG projection of n > 5 images) of the surface distribution of fluorescent PI(4,5)P_2_ (gray) and the surface distribution of 100 nM wild-type, ΔPB-Cdc11 or ΔAH-Cdc12 septin-GFP complexes (Fire LUT) at the top and bottom of cones functionalized with 2.5% mol PI(4,5)P_2_-containing membranes. Scale bar, 1 μm. **H)** Enrichment of wild-type, ΔPB-Cdc11 or ΔAH-Cdc12 mutants septin-GFP complexes at the top (κ ∼ 6.6 μm^-1^), middle (κ ∼ 2 μm^-1^) and bottom (κ ∼ 1.4 μm^-1^) of nano-cones relative to the flat membrane (κ ∼ 0 μm^-1^). Nano-cones analyzed, n = 265, 277, 342, and 580; n = 269, 279, 376 and 540; n = 266, 275, 315 and 586, for wild-type, ΔPB-Cdc11 or ΔAH-Cdc12 mutants respectively, from three technical replicates. Error bars represent s.d.; One–way ANOVA test: n.s > 0.05, *P < 0.05, ** P < 0.01, **** P < 0.0001.

Consistent with their ability to associate with membranes, GFP-tagged septin complexes containing ΔPB-Cdc11, ΔAH-Cdc12, or ΔCTE-Cdc12 also organized into septin patches on 2.5% mol PI(4,5)P_2_-containing SLBs (Figure 5B). Airyscan imaging showed a similar septin-patch organization and total surface of septin occupancy (% relative to the SLB surface) between wild-type septins and septin hetero-octamers containing the two Cdc12 mutants (Figure 5D-E). In contrast, septin complexes containing ΔPB-Cdc11 assembled into smaller patches, consistent with their reduced capacity to form longer filaments (Figure 5C). Despite this feature, the ΔPB-Cdc11 septin complexes showed a significant increase in the SLB surface occupancy, as compared to wild-type or Cdc12 mutant septin complexes (Figure 5D). Similar to their wild-type counterpart, the assembly of the ΔPB-Cdc11, ΔAH-Cdc12, or ΔCTE-Cdc12 septin complexes on 2.5% mol PI(4,5)P_2_-containing membranes led to the appearance of A647-DOPE depleted regions, which however were smaller in size as compared to wild-type septin complexes (Figure 5E). The two Cdc12 mutants showed, however, a similar average size of A647-DOPE depleted areas, suggesting that deletion of the entire CTE does not further compromise the lipid-organizing properties of septins relative to the AH partial deletion.

Septin function at the plasma membrane is usually linked to curvature-associated localizations (30, 63). Therefore, we set out to evaluate the curvature-sensing ability of septin complexes containing ΔPB-Cdc11 or ΔAH-Cdc12 on 2.5% mol PI(4,5)P_2_-containing membranes. To introduce curvature, we engineered silica substrates to display an array of conic-shaped structures with a base diameter of ∼ 1.4 µm and a height of ∼ 800 nm by soft nano-imprint lithography (soft-NIL) (Figure 5F), as we previously reported (64, 65). Silica conic-shaped substrates were functionalized with 2.5% mol PI(4,5)P_2_-containing SLBs doped with 0.2% mol fluorescent TF-TMR-PI(4,5)P_2_ (Figure 5G) and the association of wild-type and mutant GFP-tagged septin complexes at the top (radius *r* ∼ 0.15 µm), middle (*r* ∼ 0.5 µm), and bottom (*r* ∼ 0.7 µm) of the structure was characterized by Airyscan microscopy (Figure 5G-H). The enrichment (*E*) of 100 nM wild-type septin hetero-octamers at a given curvature (κ = 1/*r*) of the conic structure (*I_c_*) was obtained from the GFP-tagged septin signal (*I^s^*) normalized towards the lipid intensity (*I^l^*) relative their respective signals on the flat SLB, i.e., the surface of silica without cones 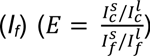. Under these experimental conditions, wild-type septins showed a preferential accumulation on positive curvatures, consistent with previous observations using lipid-functionalized silica beads and wavy substrates (40, 42). We found that wild-type septins displayed the highest accumulation at κ ∼ 6.6 μm^-1^, while this was not the case for the ΔAH-Cdc12 mutant that instead did not show preference for any positive curvatures tested. Furthermore, the ΔPB-Cdc11 only accumulated (i.e., enrichment ≥ 1.0) at κ ∼ 6.6 μm^-1^ with a relative enrichment that was ∼1.2-times lower than wild-type septins. However, the membrane binding of the ΔPB and ΔAH mutant was 4 to 3.4-times lower than that of wild-type septin complexes, respectively (Figure 5-figure supplement 2), which might explain the effect in the curvature-sensing profile of the ΔPB-Cdc11 and ΔAH-Cdc12 mutants, as curvature recognition was shown to be septin concentration–dependent (40).

### The polybasic domain of Cdc11 assists membrane remodeling by septins at the bud neck

The results above suggest that the Cdc11 PB and Cdc12 AH domains might play a role in the membrane remodeling function of septins *in cellulo*. We focused our investigations on the less explored function of the Cdc11 PB domain in cells, compared to the Cdc12 AH (41). To this end, we generated a *cdc11(Δ2-18)* mutant yeast strain that carries a deletion of the PB domain. The *cdc11(Δ2-18)* mutant was viable at 30°C, consistent with previous data (66), but temperature-sensitive at 37°C (Figure 6-figure supplement 1), indicating that the PB domain becomes important for yeast cells to survive at high temperatures. First, we monitored the recruitment of mCherry-Cdc3 to the bud neck of *cdc11(Δ2-18)* cells at 30°C by time-lapse microscopy (Figure 6A-C). We found that while the kinetics of septin recruitment in mutant cells was comparable to that of the isogenic wild-type strain, 100% of cells carrying a deletion of the PB domain failed to undergo septin ring splitting (Figure 6B). This trend was also detected at 37°C (Figure 6-figure supplement 2), where, in addition, ∼ 20% of mutant cells also displayed an absence of septin recruitment at the bud neck at the time of bud emergence (Figure 6B), thus explaining their markedly reduced viability at this temperature (Figure 6-figure supplement 1). Next, we measured septin intensity at the bud neck of *cdc11(Δ2-18)* mutant and *CDC11* wild-type small- and medium-budded cells (Figure 6C). We found a significant increase of mCherry-Cdc3 fluorescence between small- and medium-budded cells in all the conditions tested, consistent with a maturation of the septin collar during cell cycle progression. At 30°C, we observed a significant decrease in the mCherry-Cdc3 intensity at the bud neck of medium-budded *cdc11(Δ2-18)* cells compared to the *CDC11* control. This trend was, however, enhanced at 37°C, where both small- and medium budded *cdc11(Δ2-18)* mutant cells had significantly decreased septin levels at the bud neck relative to their wild type counterpart. Collectively, these results suggest that the PB domain of Cdc11 plays an important role in septin collar assembly and reorganization during the yeast cell cycle.

**Figure 6.**
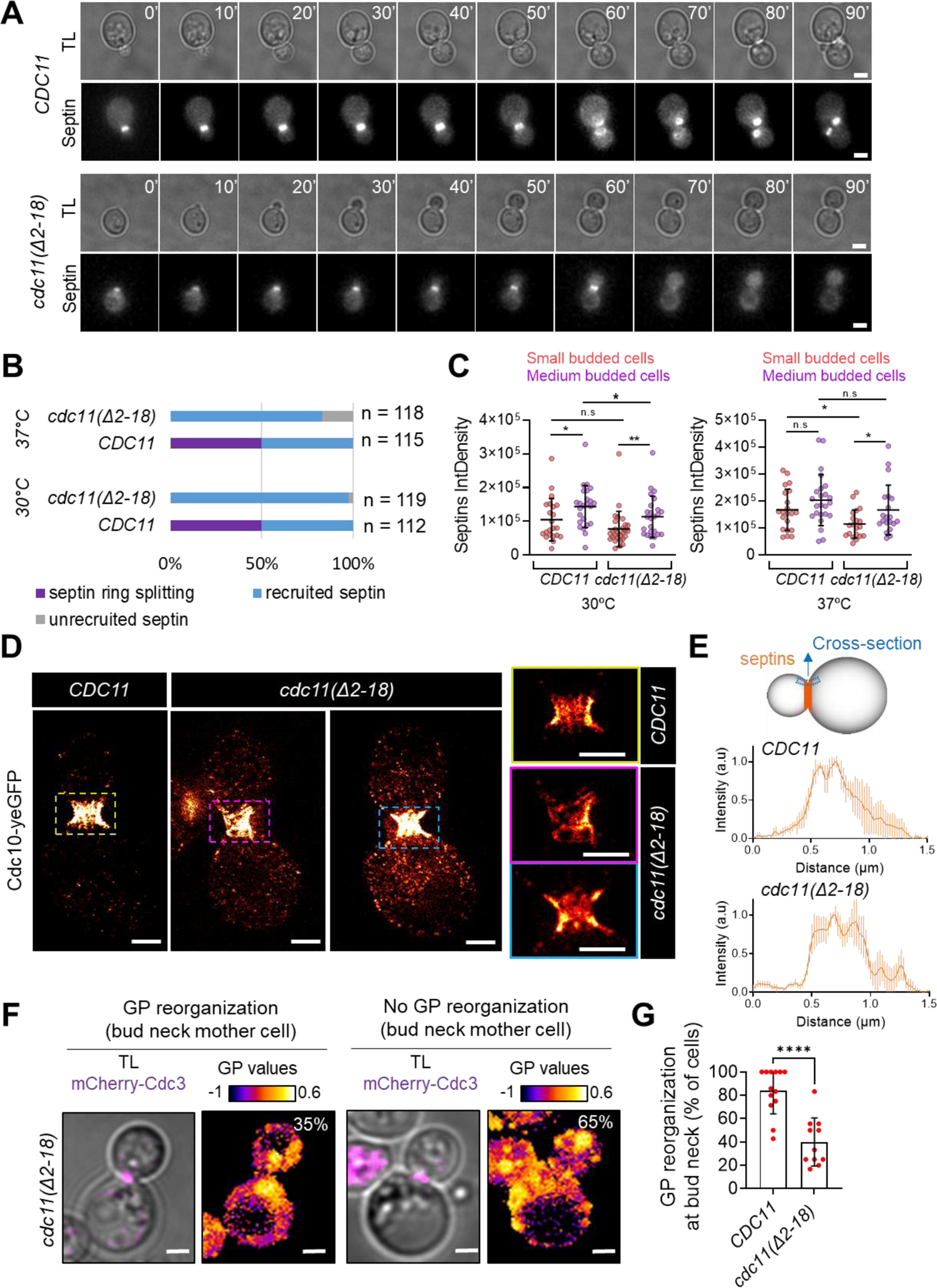
Dynamics and organization of the septin collar in wild-type and *cdc11(Δ2-18)* yeast cells. **A)** Transmitted light (TL) and TIRF images of wild-type *CDC11* and *cdc11(Δ2-18)* mutant cells expressing mCherry-Cdc3. Cells with the indicated genotypes were imaged every 5 min for 6 h at 30 °C. Scale bar, 2µm. **B)** Quantification of cells according to their phenotype for what concerns septin ring splitting (purple), unrecruited (gray), and recruited (blue) septins at the bud neck of *CDC11* and *cdc11(Δ2-18)* mutant cells expressing mCherry-Cdc3 at 30°C and 37°C. The number of cells analyzed is indicated in the graph, from two biological replicates. **C)** Septins integrated density (IntDensity) at the neck of *CDC11* and *cdc11(Δ2-18)* budding cells according to the bud size: small (orange) and medium (purple) at 30°C and 37°C. Number of structures analyzed, n = 22, n = 24, n = 28, n = 25, n = 21, n = 24, n = 20, n = 20, respectively from two biological replicates. Error bars represent s.d.; Mann-Whitney test: n.s > 0.05, * P < 0.05, ** P < 0.01. **D)** STED images of *CDC11* and *cdc11(Δ2-18)* mutant cells expressing Cdc10-yeGFP revealed with an anti-GFP nanobody Atto647N. Magnified images are identified with the corresponding color. Scale bar, 1 µm. **E)** Cross-section analysis of the outmost septin signal along the mother-bud axis as indicated in the cartoon. Average cross-section (n = 6 cells) of the anti-GFP nanobody Atto647N signal of *CDC11* and *cdc11(Δ2-18)* mutant cells. Error bars represent s.d. **F)** TL and confocal images of *cdc11(Δ2-18)* budded cells expressing mCherry-Cdc3 (magenta) and stained with the Laurdan probe. Color-coded GP value images (fire LUT) of z-stacks projections. Scale bar, 2 µm. **G)** % of *CDC11* and *cdc11(Δ2-18)* mutant cells displaying high GP values (GP > 1, indicative of Lo phases) at the bud neck of the mother cell cortex. Number of images analyzed, n = 13, and n = 11, respectively, from three biological replicates. Error bars represent s.d.; Mann-Whitney test: *** P < 0.0001.

Next, we investigated the septin collar architecture in cells lacking the PB domain of Cdc11 by STED imaging. To this end, we used wild-type *CDC11* and *cdc11(Δ2-18)* mutant cells expressing Cdc10-yeGFP that were immunolabeled with an anti-GFP nanobody Atto 647N (Figure 6D). We found that in *cdc11(Δ2-18)* cells septin structures appeared fragmented at the bud neck (Figure 6E). We also detected an increased presence of septin patches along the expected contour of the plasma membrane in mutant cells compared to the *CDC11* strain, in agreement with the pattern observed *in vitro* (Figure 5). Furthermore, the septin collar of cells lacking the PB of Cdc11 appeared less organized than the isogenic wild-type strain, which instead showed the presence of filaments ordered in parallel to the mother-bud axis, in agreement with previous reports (28, 52).

The results above indicate that the PB domain of Cdc11 is crucial for septin architecture at the bud neck of yeast cells (Figure 6B) and/or an effect of the Cdc11 PB domain on the lateral organization of the yeast cell membrane, as observed in our *in vitro* studies (Figure 5). Therefore, we set out to monitor the GP of the Laurdan probe incorporated in *CDC11* and *cdc11(Δ2-18)* mutant cells expressing mCherry-Cdc3 (Figure 6F). At 30°C, the *cdc11(Δ2-18)* mutant did not display a significant change on the global fluidity (GP index) of the plasma membrane compare to wild-type cells (Figure 6-figure supplement 3). However, we observed that the local increase in the Laurdan GP index at the mother cell bud neck overlapping with the mCherry-Cdc3 signal was significantly lower for the *cdc11(Δ2-18)* mutant cells, with only ̴ 20-30 % of cells undergoing a reorganization of high GP values (GP > 1, indicative of Lo phase), as compared to the isogenic wild-type strain (Figure 6G). To determine if other membrane-binding domains also support the effect of septins on the plasma membrane fluidity, we used the *cdc12-6* mutant that carries a C-terminal deletion that eliminates part of the AH domain (residues 390-407) (41). In the case of the *cdc12-6* mutant, experiments were performed at 25°C, as septins mostly depolymerized at 30°C (67, 68). At 25°C, cells lacking the AH domain of Cdc12 also showed a reduction in the number of cells displaying a local GP reorganization, compared to the *CDC12* wild-type strain (Figure 6-figure supplement 3). This indicates that the AH domain of Cdc12 and PB domain of Cdc11 participate in the plasma membrane-organizing function of septins in yeast cells.

## DISCUSSION

Septins are central regulators of many membrane-associated processes in eukaryotic cells (29, 30). Yet, how septins interact with and organize membranes remains a key question in the field. In this study, we investigated the function of septins as direct regulators of the lateral organization and remodeling of membranes *in cellulo* and *in vitro*. Septin collar formation at the division site of budding yeast and mammalian cells plays an essential role as a scaffold recruiting cytokinetic factors to the division site (69–71). Our results in the budding yeast *S. cerevisiae* show that localization of septins at the division site correlates with a local increase in the GP values of the environment-sensitive Laurdan probe and that this property is independent of the actin cytoskeleton. However, septins do not impact the global fluidity of the yeast plasma membrane elsewhere than at the bud neck. This observation suggests that the lipid packing effect of septins is local and linked to their membrane-bound state. Noteworthy, our results indicate that the PB and AH domains of Cdc11 and Cdc12, respectively, are required in this process.

Using STED microscopy, we reported the presence of sub-micrometric septin patches decorating the contour of the yeast plasma membrane coexisting with high-order septin structures during the budding yeast cell cycle, enlarging the repertoire of previously reported septin architectures (28, 52, 72). Our findings using a minimal reconstituted system of septin-filament assembly on fluorescently-labeled lipid membranes show that septins organization into filament-based patches appears to be regulated by the lipid composition and dependent on the functional role of the predicted membrane-interacting interfaces of yeast septins. By eliminating either the AH of Cdc11 or PB domain of Cdc12, we show that septin patch formation relies on the collective assembly of septin arrays made of at least two to five end-to-end hetero-octamers. We show that on PI(4,5)P_2_ membranes, sub-micrometric septin patches might originate from the coexistence of two septin pools. First, the membrane-bound fraction of septins might lead to the assembly of larger filament-based structures through the rotational and translational diffusion-driven interaction of octamers, consistent with previous mathematical models and molecular dynamics on rod-shaped colloidal liquids (58, 73, 74).

Indeed, the diffusive behavior of a rod-shaped structure, such as a septin octamer, might involve different states, leading to different slowing-down phenomena, especially once the local density and viscosity of the system increases. We propose that septin patch formation stems from the translational diffusive motion of septins on flat membranes, as previously reported by (75), and the existence of a subdiffusive behavior arising from a rotational, end-over-end, motion, as pointed out by the observation of a non-constant diffusion coefficient. At higher concentrations, *c* ≥1/*L*^2^, where *L* is the length of an octamer, the rotational motion of each octamer will appear severely restricted, as well as the translational motion perpendicular to the octamer axis thus, becoming the lateral interactions of short septin filaments more probable over filament annealing thus, favoring the patch appearance. The contribution of a rotational motion is also compatible with our AFM observations, showing that patches are made of short-filaments of septin arrays with different orientation angles. Second, the resulting dense septin array might promote the appearance of septin layers from the vertical or in-plane association of the solution septin fraction, consistent with previous reports on fly septins (39).

Our results show that the integrity of septin structures is particularly impaired with ΔPB-Cdc11 mutant complexes, despite their unmodified membrane-binding ability. It is possible that increased PI(4,5)P_2_ concentrations might compensate for the decreased polymerization that we observe on 2.5% mol PI(4,5)P_2_ lipid monolayers due to the PB deletion, as septin complexes carrying the same mutation are unable to polymerize in solution but form long filaments at 20% mol PI(4,5)P_2_ (17, 60).

Interestingly, septin patches have been recently found at the yeast plasma membrane under conditions that induce autophagy (72, 76). These septin patches restrain the membrane diffusion of the autophagic protein Atg9 to modulate the autophagy response. Whether the autophagy-related septin patches arise from sub-micrometric septin patches that we observe by STED remains to be investigated. Our results suggest that septin patches might fulfill two complementary roles by facilitating compartmentalized molecular interactions of septin-interacting partners and/or as reservoirs to build larger high-order septin structures. The identification of different septin architectures coexisting in cells compared to *in vitro* systems reflects the essential role of cellular factors in regulating septin assemblies, such as septin-interacting proteins(70), cytoskeletal forces(18, 25, 26), membrane topology (28), or lipid composition. In this work, we focused on the role of the lipid composition and we found that the size of septin patches grows on SLBs containing cholesterol, suggesting that in living cells, where cholesterol (or ergosterol in budding yeast) is abundant, the formation of larger septin assemblies compatible with their function as micron-scale diffusion barriers might be modulated by the lipid composition.

Septin assembly at the division site is essential for forming and stabilizing the actomyosin ring (18, 25, 26). By promoting membrane solidity through local changes in lipid packing, septin assemblies may also facilitate the stabilization of dynamic membrane remodeling processes, such as cell division (28) or cell blebbing (77), as cortex slippage can originate due to the fluid nature of the lipid membrane (78).

Using PI(4,5)P_2_-functionalized soft-NIL templates displaying micrometric to sub-micrometric positive curvatures, we showed that the PB domain of Cdc11 contributes, although to a minor extent, to the curvature-sensing properties of septins, in addition to the reported role of the AH within the CTE of Cdc12 (40, 41). Previous work showed that the PB domain of septins is important for septin polymerization and phosphoinositide specificity during membrane association (37, 45, 60). Our results with the ΔPB-Cdc11 mutant, which is largely defective in polymerization *in vitro*, suggest that septin filament assembly and membrane association is an important feature of septin localization on membranes displaying positive curvatures. Yeast cells expressing a Cdc11 mutant lacking the PB domain showed defects in septin collar architecture and lack of septin ring splitting. As stated above, this mutant also failed to engage a local membrane reorganization at the bud neck. Thus, in addition to a direct effect on septin structural organization, the PB domain of septins could indirectly impact septin architecture through their interaction with and remodeling of membranes.

Finally, our observations show that the existence of multiple curvatures in the same structure facilitates septin accumulation at sub-micrometric diameters over micrometric-scale curvatures previously observed on silica beads (40). Cooperative interactions dictate septin binding to different curvatures and depend on the sum of septin assembly parameters such as the K_on_/K_off_ rates, protein diffusion and filament annealing (43). Thus, differences in protein lateral mobility due to geometrically confined membranes (79), an enhanced PI(4,5)P_2_ interaction due to lipid defects associated with membrane curvature (80, 81) might modulate septin assembly locally.

In conclusion, we have combined *in vitro* and *in cellulo* systems to uncover the molecular mechanisms that place septins as direct regulators of the lateral organization of membranes. Cytoskeleton-driven membrane compartmentalization is essential to support cellular functions, from signal transduction to cell adhesion and migration, and this feature has been largely studied for the actin cytoskeleton (5, 8). Septins fulfill a central role as cytoskeleton scaffolds in many eukaryotic organisms and, unlike actin, directly bind membranes. Therefore, it will be important to explore further the exact functional role of septin-mediated local membrane compartmentalization in living cells.

## MATERIALS AND METHODS

### Lipids and reagents

Natural and synthetic phospholipids, including DOPC, DPPC, cholesterol, POPE, Egg-PC, Liver-PI, Brain-PI(4,5)P_2_ and fluorescent TopFluor-TMR-PI(4,5)P_2_, TopFluor-TMR-PC, Rhodamine-PE and Laurdan (6-Dodecanoyl-2-dimethylaminonaphthalene) are from Avanti Polar Lipids, Inc. Atto647N-DOPE was from Sigma. DSPE-PEG-ASP was from Abberior. Polyclonal rabbit anti-Cdc11 antibodies were from Santa Cruz (Sc-7170), DyLight 800 conjugated anti-rabbit secondary antibody was from Invitrogen (SA5-10044) and Latrunculin A was from Sigma-Aldrich.

### Yeast strains and plasmids

All yeast strains and plasmids used in this study are listed in Supplementary Table S2 and S3. Yiplac211 plasmid bearing *cdc11(Δ2-18)* was generated by Gibson assembly and verified by sequencing. Yiplac211 plasmid bearing *CDC11* or *cdc11(Δ2-18)* were integrated at the *ura3* locus of a heterozygous *CDC11/cdc11::TRP1* diploid strain after cutting with ApaI. Single copy integration was verified by Southern blot. Yeast *cdc11::TRP1* haploid strains complemented by the integrated *CDC11* or *cdc11(Δ2-18)* constructs (ySP17533 and ySP17529, respectively) were generated by sporulation and subsequent tetrad dissection. All primers used for the generation of plasmids are listed in Supplementary Table S4.

### Yeast growth conditions

Yeast cultures were grown at an appropriate temperature (i.e., 30°C for wild-type and *cdc11(Δ2-18)* strains, and 25°C for the *cdc12-6* mutant strain) overnight in synthetic SD medium containing 2% glucose (live cell imaging) or in YEPD (STED microscopy). For Laurdan experiments, the Laurdan dye was resuspended as a 5mM stock solution in DMSO and used at 5μM. Cells were incubated for 1 hour at controlled temperature under agitation (i.e., 30°C for wild-type and *cdc11(Δ2-18)* strains, and 25°C in the case of the *cdc12-6* mutant strain). Latrunculin A (LatA) was prepared as a 10mM stock solution in DMSO and cells were incubated with 0.1mM LatA for 10 minutes at appropriate temperature before Laurdan staining (i.e., 30°C for wild-type and *cdc11(Δ2-18)* strains, and 25°C in the case of the *cdc12-6* mutant strain).

### Protein expression and purification

Plasmids encoding either MBP-Cdc12/(6xHis)Cdc3-GFP or Cdc10/Cdc11 (82) were co-transformed into *E.coli* strain BL21 (DE3) Rosetta cells, then purified by amylose resin affinity and Ni-NTA-agarose beads as described (53). Cdc11-capped septin octamers containing wild-type Cdc11-Cdc12-(6His)-Cdc3-GFP-Cdc10 complex or mutants (Supplementary Table S2) were produced and purified following the same protocol. Septins were stored in storage buffer containing 20 mM Tris pH 8, 300 mM NaCl, 0.2 mM MgCl_2_, and 2 mM DTT. For *in vitro* experiments, 5 nM septin complexes were polymerized in septin polymerization buffer containing 20 mM Tris pH 8, 2 mM MgCl_2_, 400 µM GTP, 150 mM NaCl. Recombinant GST-eGFP-PH-domain (PLCδ1) was purified as described (64).

### Liposome floatation assay

Liposomes of various lipid compositions (Table S1) were prepared by mixing stocks in chloroform/methanol. Lipids were placed in a vacuum chamber for 1 h to eliminate solvent. The lipid mixtures were resuspended in 20 mM Tris-HCl pH 8, 150 mM NaCl preheated at appropriate temperature (depending on the mixture, 45°C or 56°C for phosphatidic acid-containing mixtures) to obtain 2 mM lipid suspensions. Lipid suspensions were submitted to six cycles of freezing and thawing using liquid nitrogen and a water bath at 45 °C or 56°C. Lipid suspensions were then extruded 15 times using an Avanti Mini-extruder with 100 nm polycarbonate filters (to give ∼100 nm liposomes). Proteins and liposomes (1 μM GFP-tagged septin hetero-octamers and 2 mM total lipids) were incubated 30 min in 50 μl septin buffer at room temperature in 200 µl thick-walled polycarbonate tubes (adapted to a TLA 100 Beckman rotor). The protein–lipid suspensions were adjusted to 30% w·vol^−1^ sucrose by adding and mixing 50 μl of 60% sucrose in septin buffer and overlaid with two cushions: 50 μl of 25% sucrose and 50 μl of septin buffer with no sucrose. The tubes were then centrifuged at 200,000 *g* for 1 h at 22 °C (TL 100, Beckman). After ultracentrifugation, 30 μl fractions were collected at the top and bottom and GFP-tagged septin hetero-octamers were precipitated with 10% trichloroacetic acid. The fractions were analyzed by conventional western blotting using a polyclonal rabbit anti-Cdc11 antibody and a rabbit Dylight 800. Immunodetection was performed using the Odyssey® imaging system (LI-COR Biosciences).

### Supported lipid bilayers

Glass coverslips were cleaned by sequential 30 min sonication steps in 1 M NaOH,100% ethanol and, milliQ water, with water rinsing between each step. The lipid mixtures of supported lipid bilayers (SLBs) consisted of: 70-80% Egg-PC, 20-25% Liver-PI and 2.5-5% of Brain-PI(4,5)P_2_. In the case of the ternary mixture, the lipid composition of SLBs was: mica-supported lipid membranes made of 2.5% mol Brain-PI(4,5)P_2_, 2.5% mol DOPC, 30% mol DPPC, 44.8% mol cholesterol, 20% mol Liver-PI. The amount of total negatively charged lipids was kept to 25% for all the mixtures containing phosphoinositides at the expenses of Liver-PI. Fluorescent lipid analogs were added at 0.2% mol.

For fluorescence microscopy and AFM experiments, supported lipid bilayers were prepared as follows. Briefly, lipids dissolved in methanol/chloroform were mixed and dried for 1h in a vacuum oven at 60°C. Small unilamellar vesicles (SUVs, diameter ∼ 100nm) were obtained by extrusion of multilamellar vesicles in citrate buffer (citrate 20 mM pH 4.6, KCl 50 mM, EGTA 0.5 mM) heated to 44°C. 40 µl of SUVs were deposited on a two-well silicone insert (IBIDI) and incubated 40 min at 40°C. Bilayers were carefully rinsed with wash buffer (20 mM Tris-HCl pH 8, 300 mM NaCl) and then equilibrated with equilibration buffer (20 mM Tris-HCl pH 8, 150 mM NaCl, 1 mg/ml lipid-free BSA). GFP-tagged septin hetero-octamers (wild-type or mutants) were diluted to 10 nM in septin buffer (20 mM Tris-HCl pH 8, 150 mM NaCl, 2 mM MgCl_2_, 400 µM GTP). Septins complexes at a final concentration of 5 nM were added on SLBs.

### Silica thin film nanostructuration

The master used for the nano-imprint lithography is made of undoped silicon (100) 300 µm thick. First, the substrate is cleaned with Piranha. Then a 100 nm chromium layer is deposited by sputtering (UNIVEX 150), next a negative photoresist (Az2020) is spin-coated on the substrate and a UV lithography is performed through a mask. The mask’s design is a periodic nano-holes pattern made in a chromium layer, the nano-holes diameters are 2 µm wide with a pitch of 4 µm. After development (1min in AZ 726) the obtained pattern is an array of nano-disks, with a 2 µm diameter and a 4 µm pitch, of photoresist. These disks are then thinned with an oxygen plasma generated by an Inductively Coupled Plasma – Reactive Ion Etching equipment (ICP RIE Corial 150L) in order to achieve a 700 nm diameter. After the ICP-RIE process, the chromium layer is etched with a chromium-based solution (Chromium etchant standard from Sigma Aldrich 651826) for 5 min revealing a nano-disks pattern (the negative pattern of the mask) on the chromium. These disks are used as hardmask for the plasma etching of the Si substrate. The RIE recipe use CHF3 plasma as etchant and the conditions (Power, Pressure, Flow) are defined to obtain a troncoïdal shape for the pillars. The last step is the sample cleaning with Piranha.

Next, SiO_2_ vertical cone-shape nanostructures were prepared on conventional borosilicate coverslips with precision thickness No. 1.5 (0.170 ± 0.005 mm) by soft nano-imprint lithography (soft-NIL), as previously reported (64, 65). The dimensions of the silicon master and the resulting printed silica nanostructures were characterized by scanning electron microscopy (SEM).

### Airyscan/confocal microscopy and fluorescence recovery after photobleaching imaging

SLBs and septins were imaged on a Zeiss LSM880 Airyscan/confocal microscope. Excitation sources used were: 405 nm diode laser, an Argon laser for 488 nm and 514 nm and a Helium/Neon laser for 633 nm. Acquisitions were performed on a 63x/1.4 objective.

Multidimensional acquisitions were acquired via an Airyscan detector (32-channel GaAsP photomultiplier tube (PMT) array detector). For fluorescence recovery after photobleaching (FRAP) measurements, GFP-tagged septin hetero-octamers were photobleached over a circular or square area using 50 iterations of the 488 nm laser with 70% laser power transmission. We recorded n =20 pre-bleaching images every 0.042 s or for long acquisitions (i.e. ≥ 7 min), every 30 s.

The half-life (t_1/2_), which is the time at the half-maximal fluorescence intensity from the plateau, and the immobile (*IM_f_*) and mobile fraction (*M_f_*) of septins bound to membranes were estimated after normalization of fluorescence intensities and exponential fitting of FRAP curves using Prism GraphPad software. We obtained the % of *M_f_* and *IM_f_* by applying the following equations: *M_f_ = I_∞_ − I_0_/I_i_ − I_0_*, where *I*_∞_ is the fluorescence intensity at the plateau, *I*_0_ is the minimal intensity after bleaching, and *I*_i_ is the maximal intensity before bleaching.

### Atomic Force Microscopy

AFM experiments were performed at room temperature on a JPK NanoWizard IV XP microscope (JPK BioAFM, Bruker Nano GmBH, Berlin, Germany) mounted on an inverted Nikon Ti2-U microscope (Nikon Instruments Europe B.V., Amsterdam, The Netherlands) equipped with a standard monochrome CCD camera (ProgRes MFCool, Jenoptik, Jena, Germany). Fluorescence imaging was performed by wide-field illumination (Intensilight Hg lamp, Nikon) with a 100X objective (Nikon CFI Apo VC, 1.4 NA, oil immersion) and the appropriate filter cubes. A software module (DirectOverlay, JPK Bio-AFM, Bruker Nano GmbH, Berlin, Germany) was used to calibrate the tip position with the optical image. AFM images were acquired in Quantitative Imaging (QI™) mode under a setpoint force of 250 pN, based on the previous works (83). Rectangular-shaped BL-AC40TS-C2 cantilevers (Olympus) with a nominal spring constant of 0.09 N·m^-1^ and a nominal tip radius of 8 nm were used for septin nano-domain characterization and PEAKFORCE-HIRS-F-A cantilevers (Bruker) with a nominal spring constant of 0.35 N·m^-1^ and a nominal tip radius of 1 nm were used for high resolution imaging. All AFM experiments were operated under liquid conditions. Before each experiment, the cantilever spring constant and sensitivity were determined by the thermal noise method (84). QI mapping was performed at 256 x 256 pixels/line at a scan rate of 9.4 kHz. SLBs were prepared on 1M NaOH treated coverslips following the same experimental procedure detailed in the previous section. After overnight incubation of 5 nM of Septin-GFP filaments, samples were mounted on the AFM imaging holder and imaged in 20 mM Tris-HCl pH 8, 2 mM MgCl_2_, 150 mM NaCl.

### Transmission Electron Microscopy

Samples for transmission electron microscopy (TEM) were prepared by incubating GFP-tagged septin hetero-octamers (wild-type or mutants) with lipid monolayers formed at an air-buffer interface. The lipid solution consisted of 77.5% Egg-PC, 20% Liver-PI and 2.5% of Brain-PI(4,5)P_2_ in chloroform. Briefly, each Teflon well was filled with 20 µl of a solution of 0.005 mg/ml GFP-tagged septin hetero-octamers in septin buffer (20 mM Tris-HCl pH 8, 2 mM MgCl_2_, 400 µM GTP, 150 mM NaCl). A drop (∼ 0.5 µl) of a 0.5 g/L lipid solution in chloroform was deposited at the air-water interface in each well to form a lipid monolayer. The Teflon block was kept overnight at 4 °C in a humid chamber. Lipid monolayers with adsorbed GFP-tagged septin hetero-octamers were collected by briefly placing hydrophobic grids (Electron Microscopy Sciences, Hatfield, PA, USA) on the surface of the solutions with the carbon-coated side facing the solution. The lipid monolayers were absorbed on a carbon-coated grid on a Formvar carbon-coated grid (Delta Microscopies FCF100-Cu) for 30 sec. The grids were then negatively stained for 1 min using 2% uranyl acetate in water. Data were collected using a Tecnai F20 transmission electron microscope at 120KV and equipped with a Veleta camera.

### Live cell imaging

Cells were mounted on 1% agarose pads in SD medium on Fluorodishes (FD35-100 World Precision Instrument) and filmed at controlled temperatures (25 or 30°C) with a 63X 1.4 NA Oil DIC M27 immersion objective mounted on a Zeiss LSM 980 NLO microscope equipped with Chameleon Ultra-II Coherent LASER (680-1080nm). Excitation sources used were an Argon laser for 514 nm and Chameleon laser for 800 nm. Multidimensional acquisitions were acquired via GaAsP and Multialkali photomultiplier tube (PMT) array detectors. Z-stacks of 5 planes were acquired with a step size of 0.4 µm. Z stacks were max-projected with ImageJ. Selected cells were illuminated with an 800 nm laser and emission read at 420-460 nm (representing the liquid ordered phase) and 490-540 nm (representing the liquid disordered phase). For Laurdan experiments, the Laurdan dye was used at 5 μM. Cells were incubated for 1 hour at appropriate temperature under agitation. Latrunculin A (LatA) was used at 0.1 mM for 10 minutes at appropriate temperature.

### STED imaging

Samples for STED microscopy were prepared by incubating yeGFP-tagged Cdc3 cells (wild-type CDC11 or mutants) with a 1/250 dilution of an anti-GFP nanobody Atto 647N (ChromoTek), as previously detailed in (52). Untagged yeGFP yeast cells were used as a control for the nanobody specificity and background evaluation. After 90min of antibody incubation, cells were washed twice in PBS by gently shaking at room temperature and mounted using Abberior Mount Liquid (Abberior).

Images were acquired on an Expert Line STED microscope (Abberior Instruments GmbH) using a PlanSuperApo 100X oil immersion objective N.A. 1.4 (Olympus). Atto 647N was imaged at 640 nm excitation using a 775nm depletion laser line. Detection was set to 650-750nm. A dwell time of 10µs was used. Images were collected in line accumulation mode (5 lines), the pinhole was set to 1.0 Airy units and a pixel size of 20nm was used for all acquisitions.

### Image processing and quantification

Line profiles of the fluorescence intensities and the kymographs were made using ImageJ. The semi-automatic analysis of images to determine DOPE-depleted area and septin domains size were done using the macro toolset in ImageJ. Quantifications of septin binding were done using ImageJ. Septin filaments size was measured using FiberApp (85).

AFM image processing and profile analysis were performed using the open-source software Gwyddion (86). Membrane stiffness (Young modulus) was obtained based on the Hertzian model for elastic indentations. Young’s moduli were calculated at a sample deformations that was < 20% of the bilayer thickness (87), as previously reported (83). Correlative fluorescence and AFM image representation were performed using ImageJ (88) and the BigWarp plugin (89).

Generalized Polarization (GP) values were calculated for each pixel of yeast plasma membrane using a GP calculation macro toolset according to the following equation: GP = I_420_ −I_490_ / I_420_ +I_490_ where I_420_ and I_490_ represent the intensity of pixels in the areas of interest in the image acquired in the ordered and disordered spectral channels, respectively. The cross-sections showing the GP values at the yeast plasma membrane were done with ImageJ.

### Data representation and statistical analysis

Representation of cross-section analysis and recruitment curves were performed using Origin and Prism GraphPad software. Statistical analysis was performed using the Mann-Whitney two-tailed test or ordinary one-way ANOVA multiple comparisons test, with single pooled variance using Prism GraphPad software. In all statistical, the levels of significance were defined as: *P<0.05, **P<0.01, ***P<0.001 and ****P<0.0001.

## Data availability

All data generated or analyzed during this study are included in the manuscript and supporting file; source data files have been provided.

## ACKNOWLEDGMENTS

We acknowledge Erdinc Sezgin for help with the Lo marker DSPE-PEG-ASR. We also thank Julien Viaud for kindly providing the GFP-PH^PLCδ^construct. The authors are grateful to Alenka Copic and Cyril Favard for helpful discussions. This work has been supported by a grant of the Agence Nationale pour la Recherche (SEPTORG ANR-18-CE13-0015-01) to S.P. and L.P. We acknowledge the imaging facility MRI, member of the national infrastructure France-BioImaging supported by the French National Research Agency (ANR-10-INBS-04, «Investments for the future»).

## AUTHOR CONTRIBUTIONS

Conceptualization: L.P, S.P. and F.E.A. Methodology: F.E.A., S.I., S.L., C.C., D.S, R.D, M-P.B, A.P, S.P. and L.P. Supervision: A.C-G, L.P. and S.P. Funding acquisition: L.P. and S.P. Writing: L.P and F.E.A. with inputs from authors.

## SUPPLEMENTARY INFORMATION

**Table S1.** Composition of the different SUVs lipid mixtures used for the liposome floatation assay.

|  | SUVs mixture |  |
| --- | --- | --- |
|  | Lipid | Mole % |
| Compositon PI(4,5)P<sub>2</sub> | Egg-PC | 80 |
|  | Brain-PI(4,5)P<sub>2</sub> | 5 |
|  | Liver-PI | 15 |
| Compositon PI | Egg-PC | 75 |
|  | Liver-PI | 25 |
| Compositon PA | Egg-PC | 75 |
|  | Egg-PA | 10 |
|  | Liver-PI | 15 |
| Compositon PS | Egg-PC | 75 |
|  | Brain-PS | 10 |
|  | Liver-PI | 15 |
| Compositon PC | Egg-PC | 100 |

**Table S2.** List of *S. cerevisiae* strains used in this study.

| Name | Relevant genotype |
| --- | --- |
| ySP41 | *MATa, ade2-1, trp1-1, can1-100, leu2-3,112, his3-11,15, ura3* |
| ySP15355 | *MATalpha, ade2-1, trp1-1, can1-100, leu2-3,112, his3-11,15, ura3, bud4::LEU2:: BUD4, cdc3::mCherry-CDC3::URA3* |
| ySP16736 | *MATa, bud4::LEU2::BUD4, cdc3::mCherry-CDC3::URA3* |
| ySP17147 | *MATa, cdc3::mCherry-CDC3::URA3, cdc12-6* |
| ySP17529 | *MATalpha, cdc11::TRP1, ura3:URA3::cdc11(Δ2-18), bud4::LEU2::BUD4, cdc3::mCherry-CDC3::URA3* |
| ySP17533 | *MATalpha, cdc11::TRP1, ura3:URA3::CDC11, bud4::LEU2::BUD4, cdc3::mCherry-CDC3::URA3* |
| ySP17816 | *MATa, cdc11::TRP1, ura3:URA3::CDC11, bud4::LEU2::BUD4, CDC10-yeGFP::KanMX* |
| ySP17818 | *MATa, cdc11::TRP1, ura3:URA3::cdc11(Δ2-18), bud4::LEU2::BUD4, CDC10-yeGFP::KanMX* |

**Table S3.**
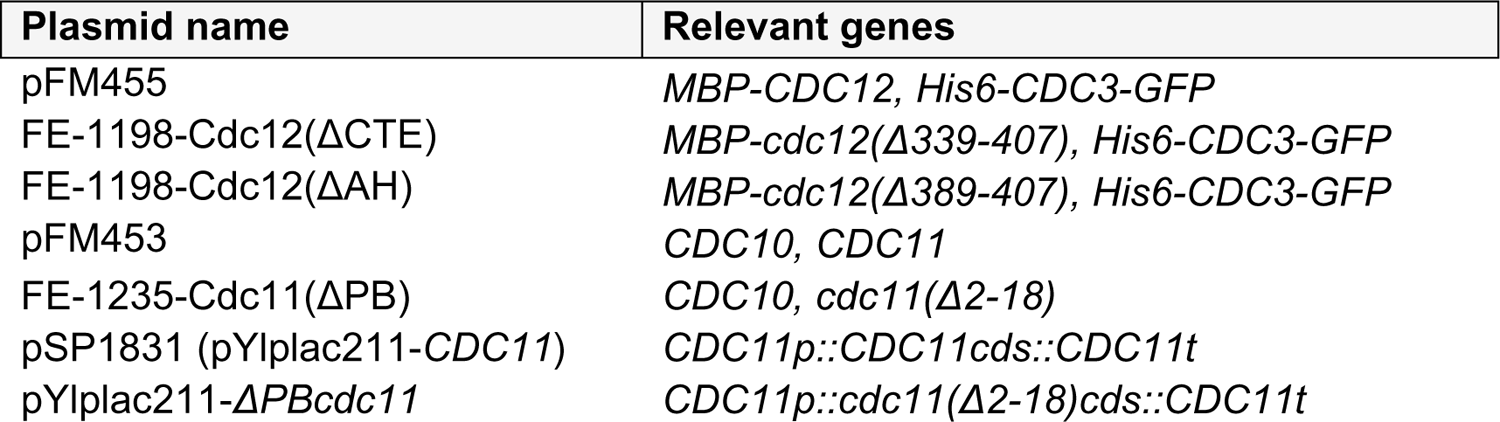
List of plasmids used in this study.

**Table S4.**
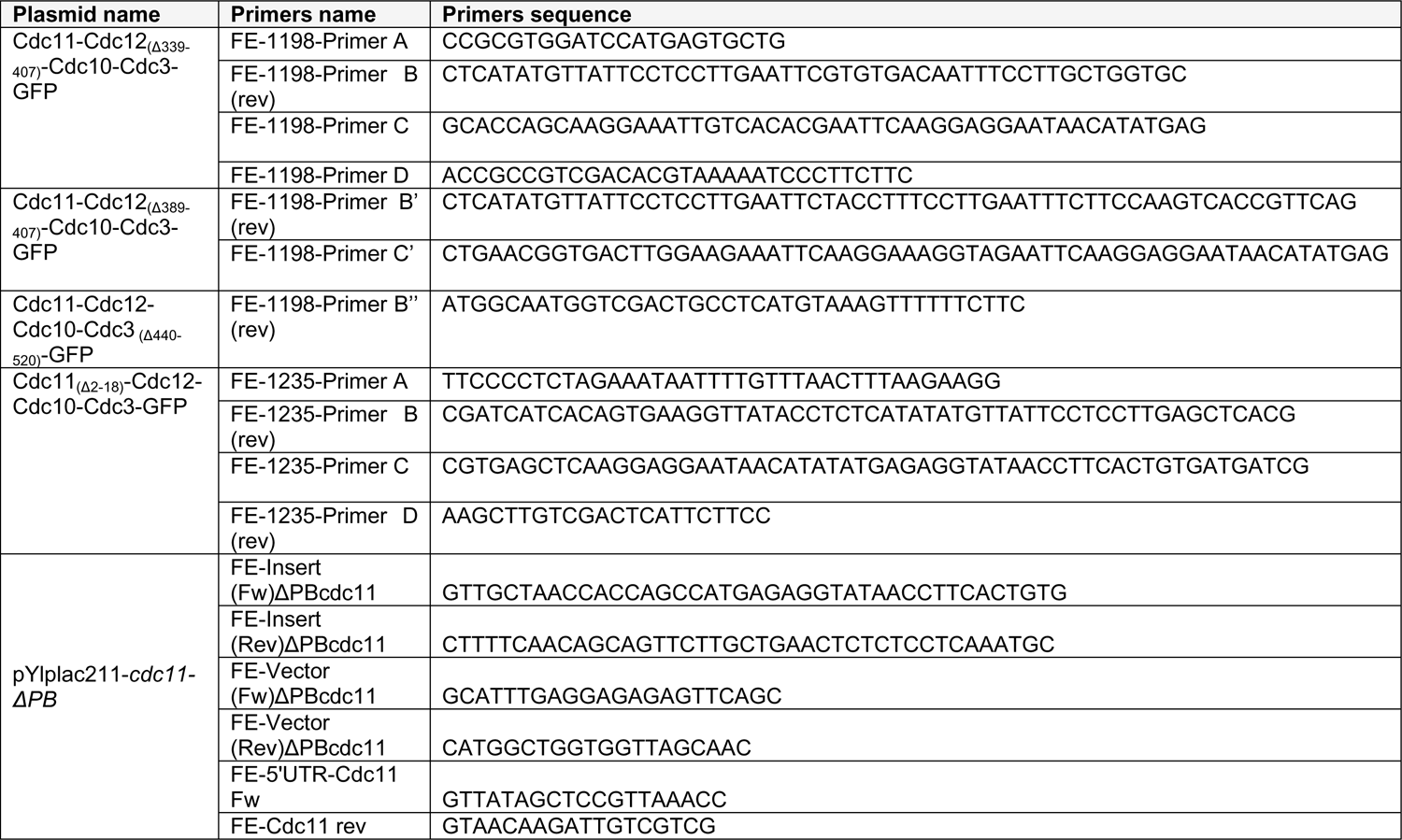
List of primers used in this study.

**Figure 1-figure supplement 1.**
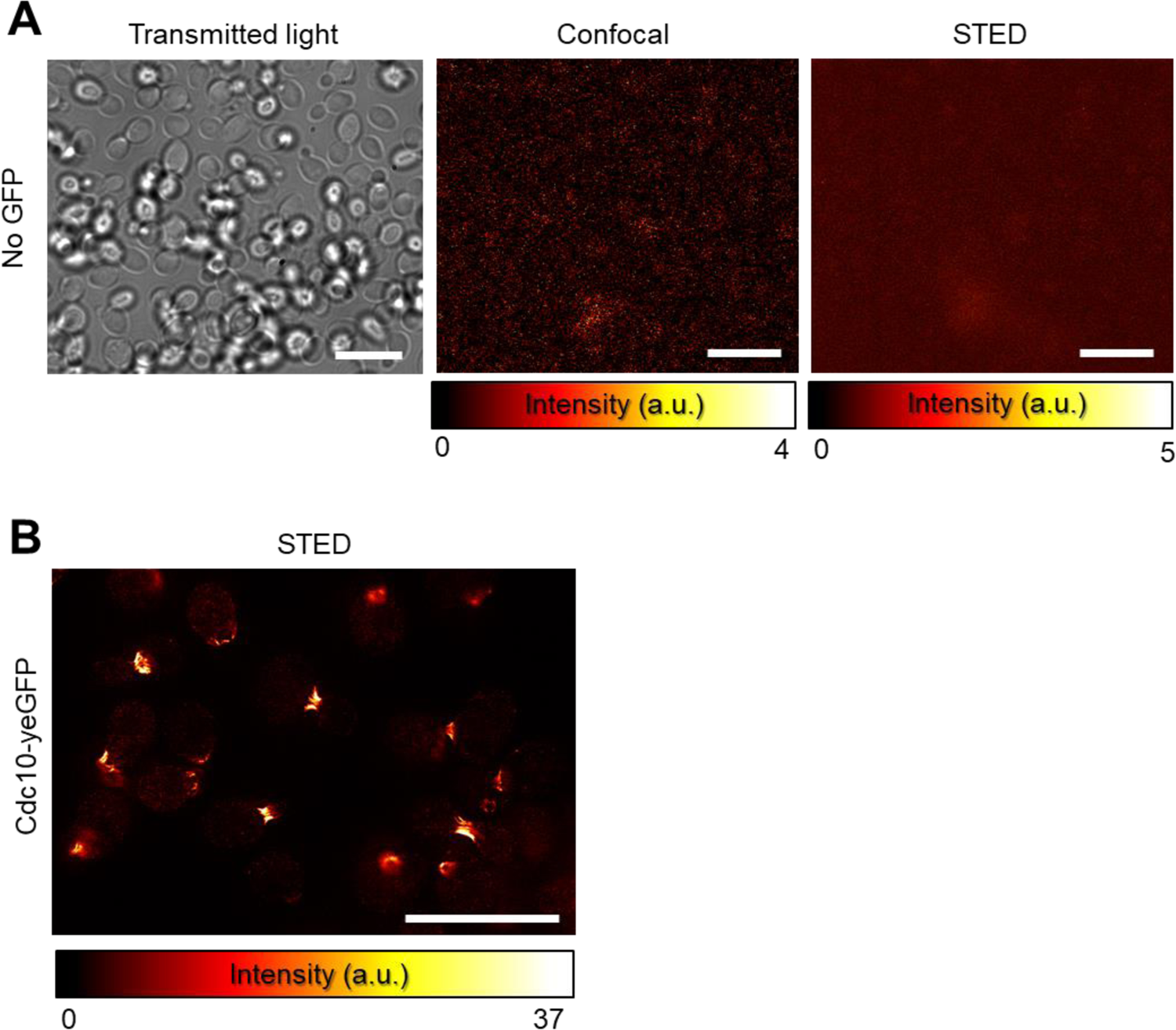
**A-B)** Transmitted light, confocal and STED images of wild-type yeast cells (i.e., no expressing yeGFP-tagged proteins) **(A)** compared to a STED image of yeast cells expressing Cdc10-yeGFP **(B)**. All conditions were immunostained with an anti-GFP nanobody Atto647N. Images were acquired under the same experimental settings. Gray scale of confocal and STED images is displayed in a Fire LUT. Scale bar, 10 µm.

**Figure 2-figure supplement 1.**
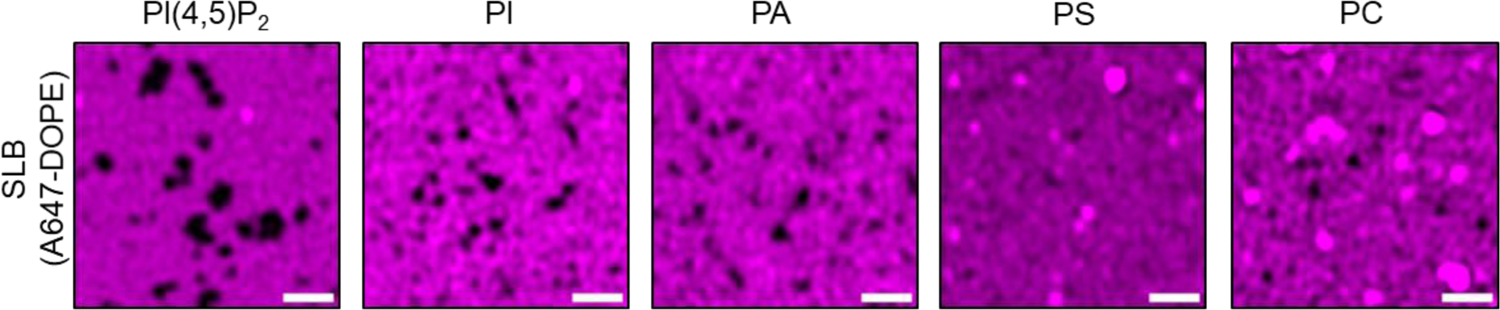
Airyscan images showing the signal of fluorescent A647-DOPE (0.2% mol) on lipid bilayers (SLBs) made with the SUVs compositions detailed in Table S1 after addition of 5 nM septin-GFP hetero-octamers prepared in septins buffer (150 mM NaCl). Scale bar, 5 µm.

**Figure 3-figure supplement 1.**
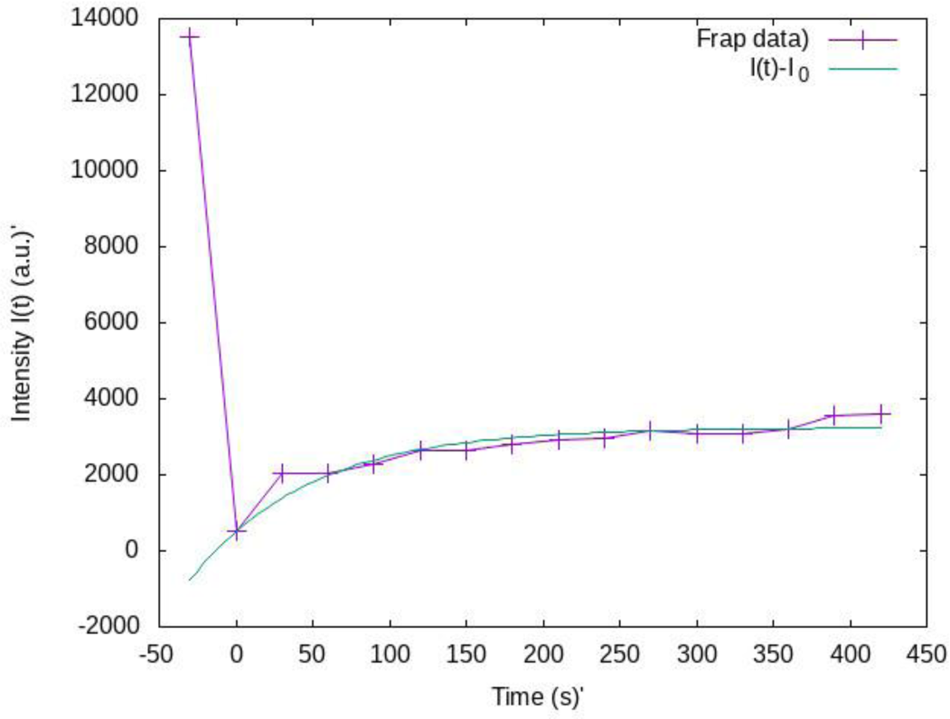
Fitting of the FRAP analysis. The fitting function, based on Brownian motion theory, is: *I*(*t*) − *I*_0_ = *I*_∞_ (1 − *e*^−*t*⁄τ^) and gives *I* = (2725 ± 110) (*a. u*); ^1^⁄_τ_ = (0.013 ± 0.002) *S*^−1^ or also τ = (77 ± 12)*S* for *I*_0_ = 515 (*a. u*.). The time τ is the typical recovery time of the fluorescent signal. We can then estimate the mobile fraction 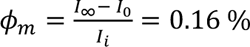 and the immobile fraction Φ_*imm*_ = 1 − Φ_*m*_ = 0.84 %. We can associate an effective diffusion constant of the septin octamers in the patch (during the experimental time) to the relaxation time τ, indeed,

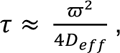

where ω = 1,75 μ*m* is the typical radius of the photobleached spot:

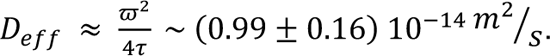

**Figure 3-figure supplement 2.**
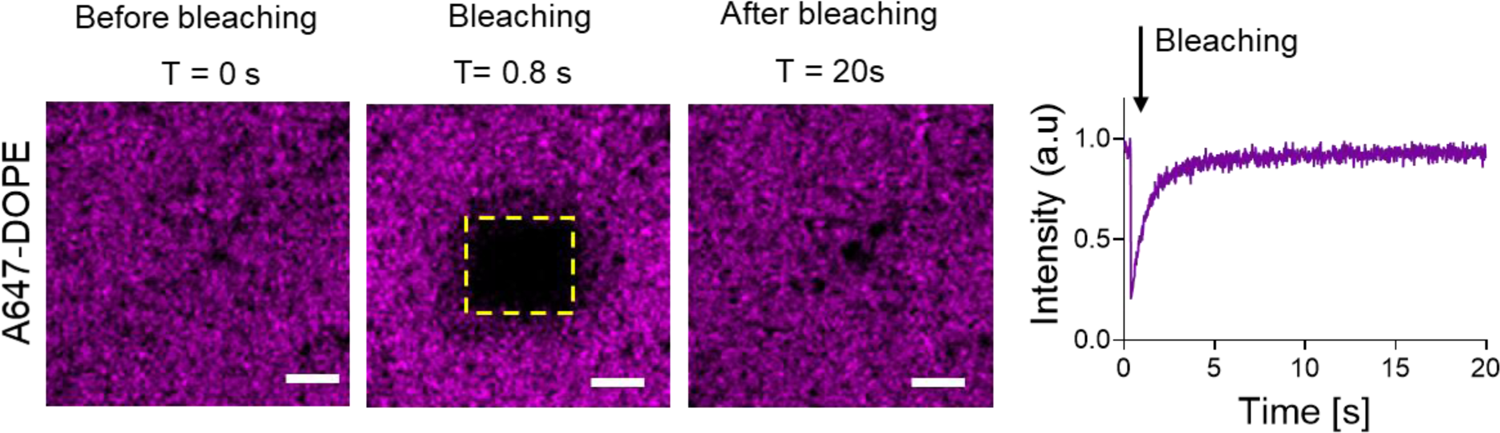
Fluorescence recovery of the A647-DOPE lipid signal of SLBs made of 77.3% mol Egg-PC, 20% mol Liver-PI; 2.5% mol Brain-PI(4,5)P2, and 0.2% mol fluorescent A647-DOPE and incubated with 5 nM of septins-GFP. The FRAP experiment was performed on yellow ROI indicated. Snapshots of the SLBs corresponding to the A647-DOPE channel before, at the moment of bleaching and after bleaching. Representation of the fluorescence intensity of A647-DOPE over time acquired at the FRAP ROI. Scale bar, 1 μm.

**Figure 3-figure supplement 3.**
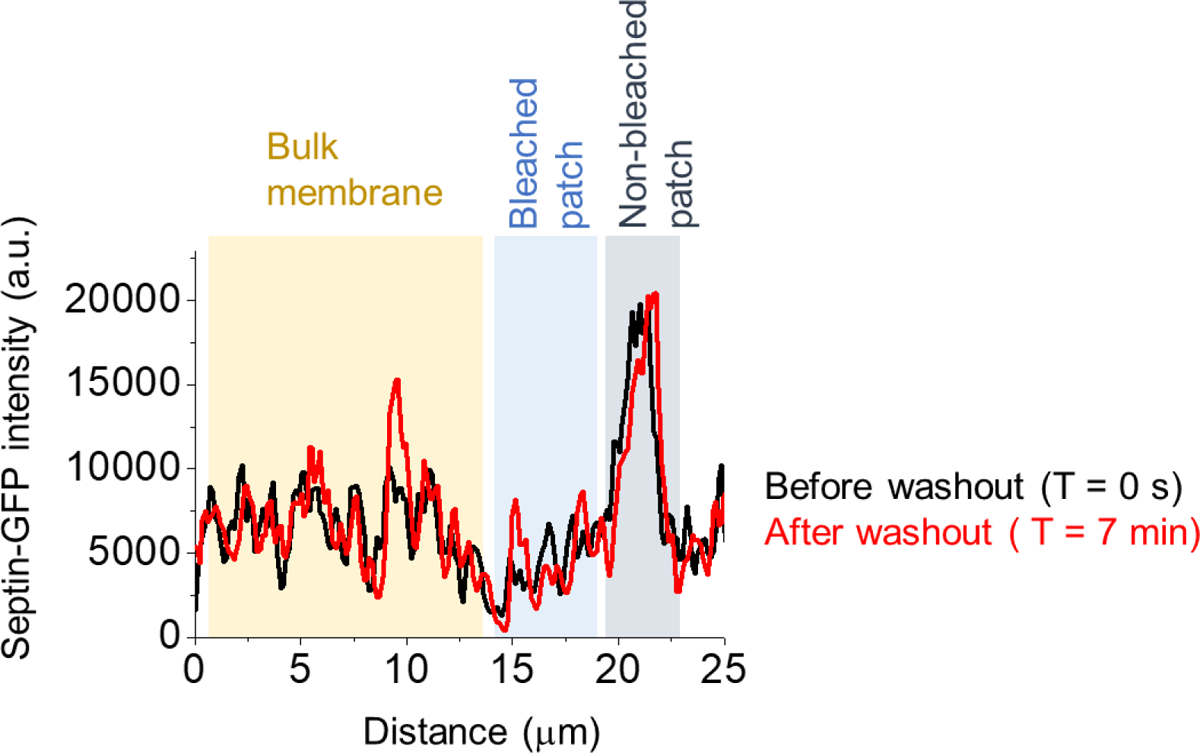
Cross-section of the septin-GFP signal along the orange line in Figure 3A after the bleaching process (T = 420s) before (black) and after (red) septin wash out. The different features of the septin signal on SLBs are highlighted (septins at the bulk membrane and bleached and non-bleached septin patch).

**Figure 4-figure supplement 1.**
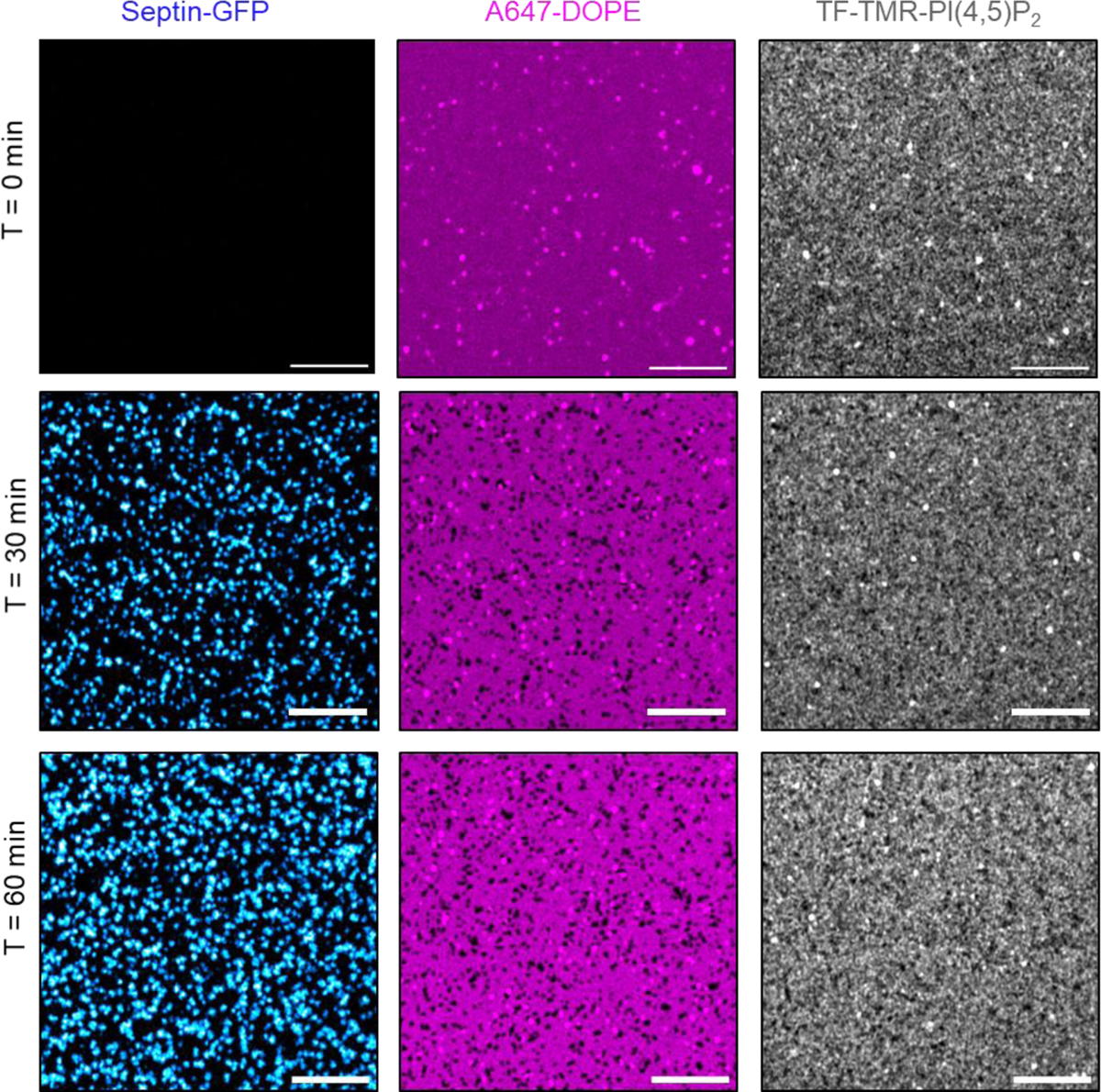
Time-dependent effect on the formation of DOPE-depleted domains by septin nano-domains upon binding on PI(4,5)P_2_-containing membranes. Airyscan images showing septin association with lipid bilayers containing 77.4% mol Egg-PC, 20% mol Liver-PI; 2.4% mol Brain-PI(4,5)P2, 0.2% mol fluorescent TF-TMR-PI(4,5)P_2_ and 0.2% mol fluorescent A647-DOPE, at time T = 0 min, 30 min and 60 min. 5 nM septin-GFP hetero-octamers were added in septin buffer (150 mM NaCl). Scale bar, 5 μm.

**Figure 4-figure supplement 2.**
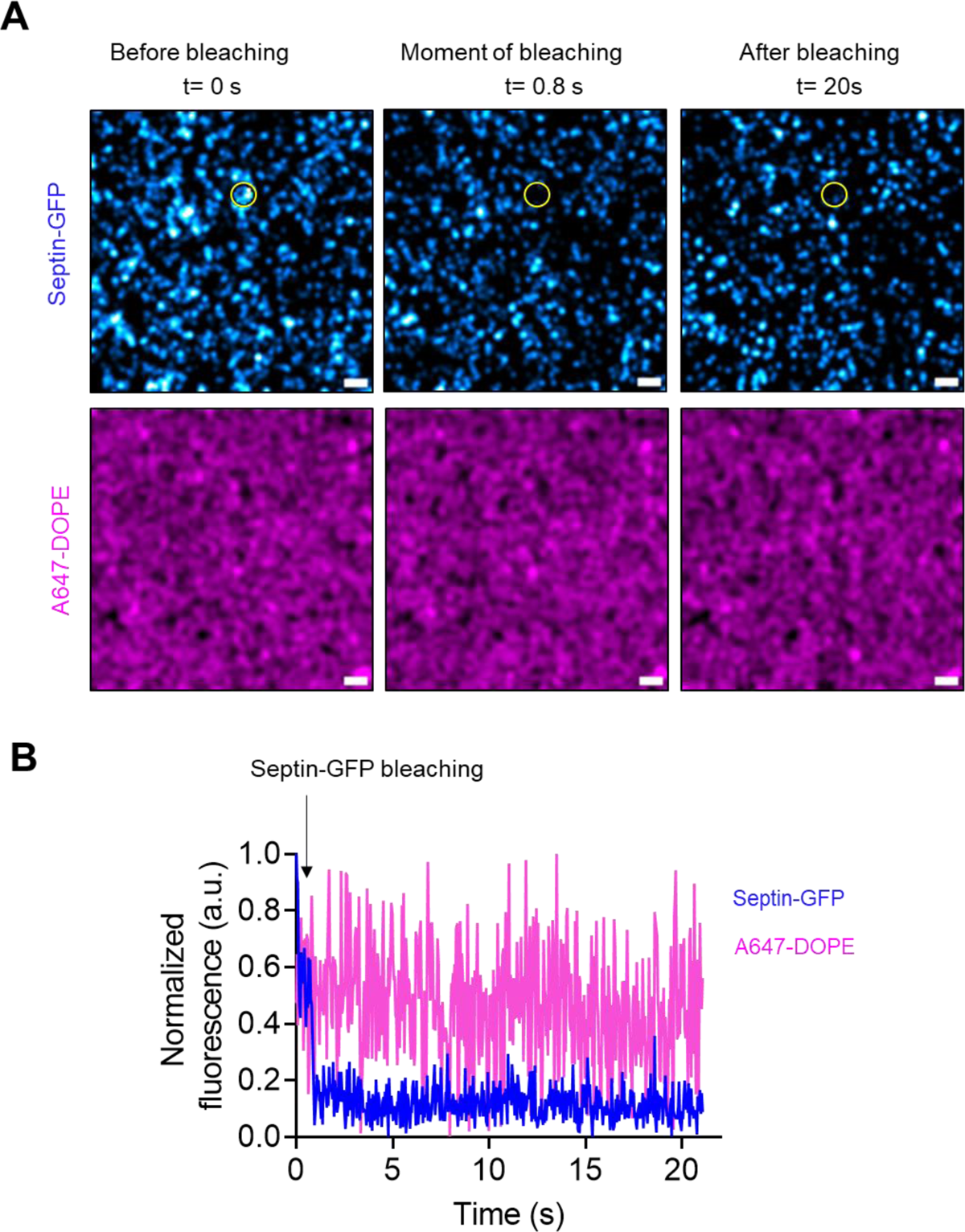
Effect of septin-GFP bleaching on the A647-DOPE lipid signal. The FRAP experiment was performed on the septins-GFP channel after septins were let for 10 min to bind SLBs containing 77.3% mol Egg-PC, 20% mol Liver-PI; 2.5% mol Brain-PI(4,5)P2, and 0.2% mol fluorescent A647-DOPE. **A)** Snapshots of the septin-GFP and A647-DOPE channel before, at the moment of bleaching and after bleaching a ROI of ∼ 1 μm diameter on the septin-GFP channel and corresponding to a septin nano-domain. **B)** Fluorescence intensity of septin-GFP and A647-DOPE over time acquired at the FRAP ROI. Scale bar, 1 μm.

**Figure 4-figure supplement 3.**
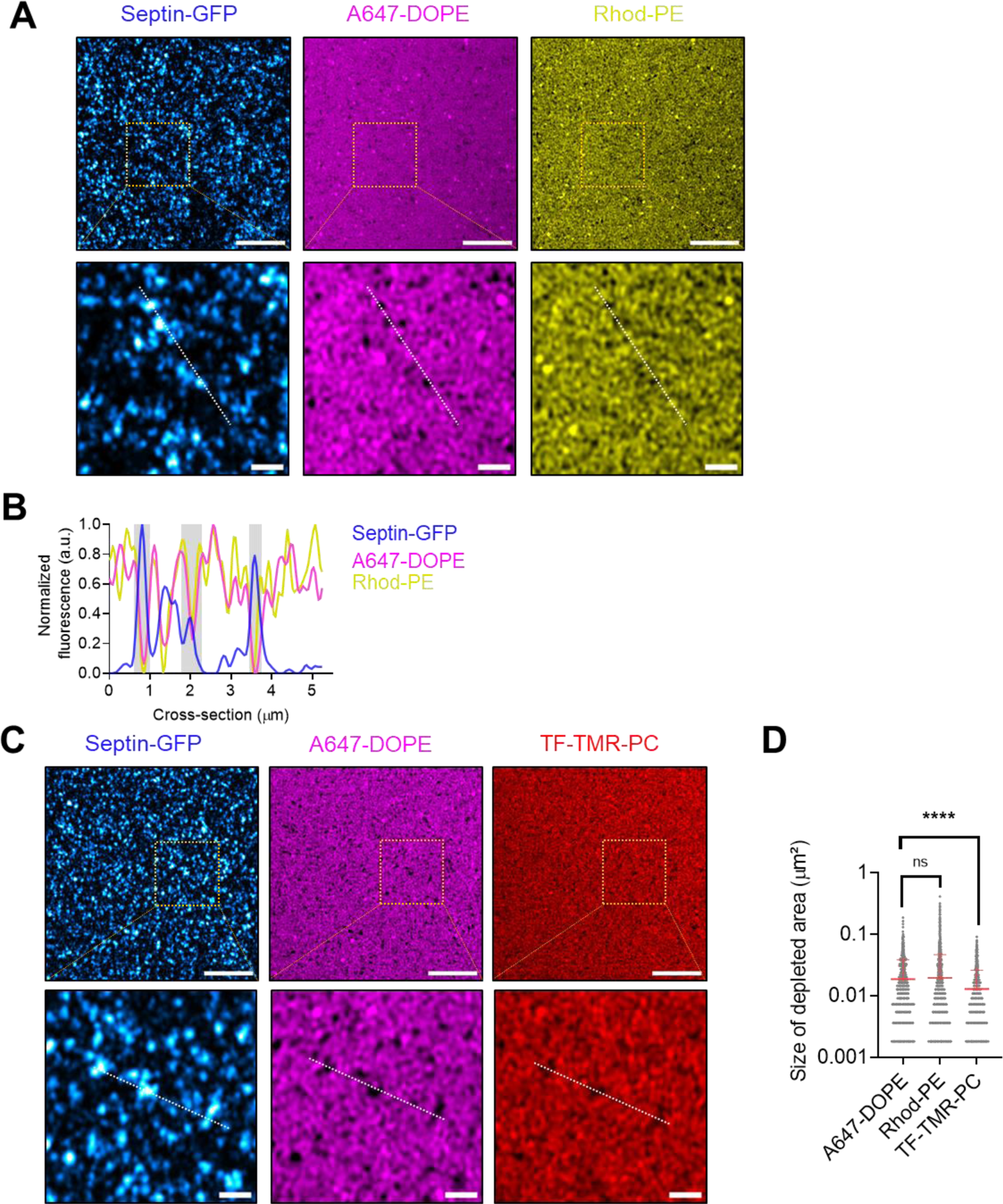
**A)** Airyscan images and **B)** cross-section along the red-dashed line showing the septin-GFP hetero-octamers on 2.5% mol Brain-PI(4,5)P_2_-containing SLBs doped with 0.2% mol fluorescent A647-DOPE and 0.2% mol fluorescent Rhodamine-DOPE. **C)** Airyscan images showing the septins-GFP hetero-octamers on 2.5% mol Brain-PI(4,5)P_2_-containing SLBs doped with 0.2% mol fluorescent A647-DOPE and 0.2% mol fluorescent TF-TMR-PC. **D)** Size of lipid-depleted areas calculated on Airyscan images. The number of depleted areas measured for A647-DOPE, Rhodamine-DOPE, and TF-TMR-PC are n = 1024, n = 3749 and n= 1448, respectively. Scale bar, 5 μm.

**Figure 4-figure supplement 4.**
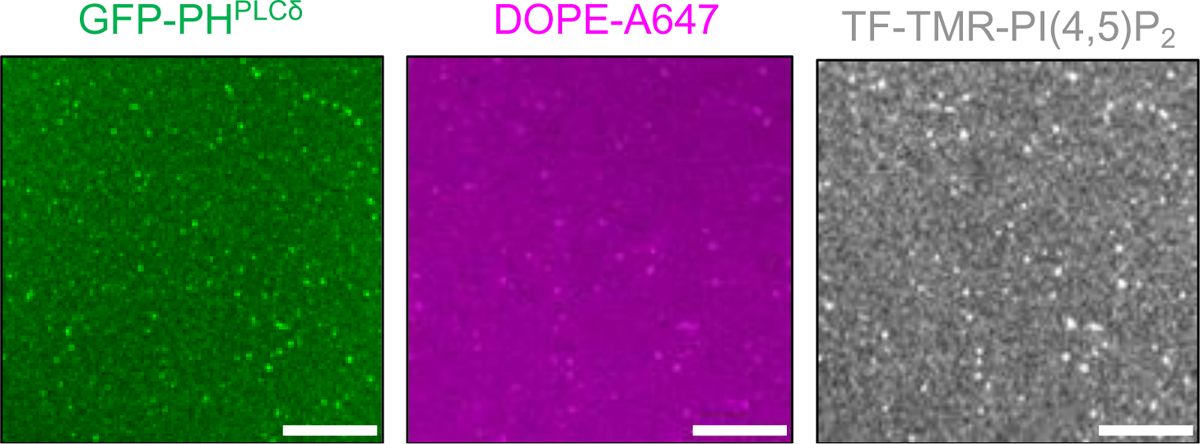
Airyscan images showing the binding of 1 µM of GFP-PH^PLCδ^ with lipid bilayers containing 77.3% mol Egg-PC, 20% mol Liver-PI; 2.3% mol Brain-PI(4,5)P_2_, 0.2% mol fluorescent TF-TMR-PI(4,5)P_2_ and 0.2% mol fluorescent DOPE-A647. Scale bar, 5 µm.

**Figure 4-figure supplement 5.**
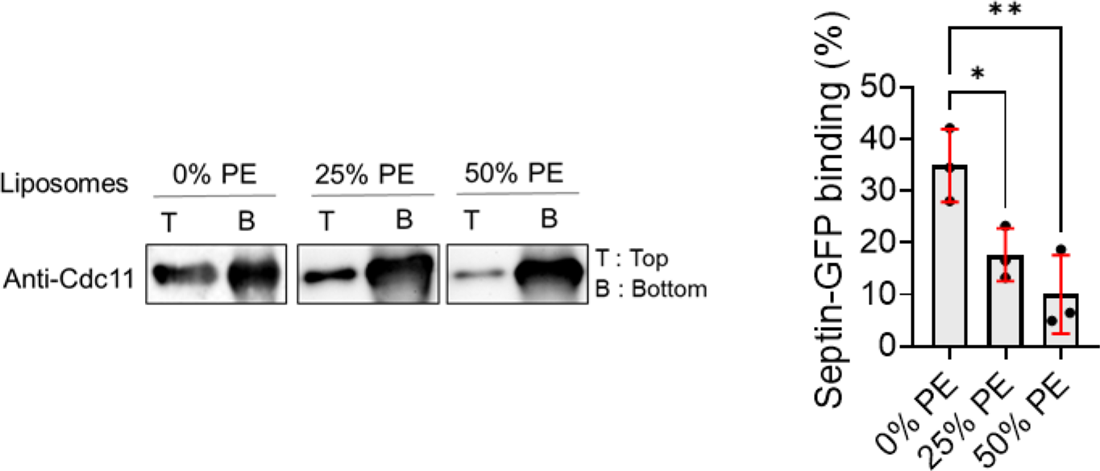
1 µM of septin-GFP hetero-octamers were incubated with liposomes (2 mM) made of Egg-PC, Liver-PI, Brain-PI(4,5)P_2_ and different % mole of 1-palmitoyl-2-oleoyl-sn-glycero-3-phosphoethanolamine (PE) (0% PE, 25% PE, or 50% PE) with 25% total negative charge. The suspension was subjected to liposome flotation assays. Septin-GFP hetero-octamers were polymerized in septin polymerization buffer and the septin Cdc11 was detected by western blotting. Quantification of septin binding to liposomes: Top/(Top+Bottom)x100. Results are presented as mean ± s.d. (n = 3). Error bars represent s.d.; Mann-Whitney test: * P < 0.05, ** P < 0.01.

**Figure 5-figure supplement 1.**
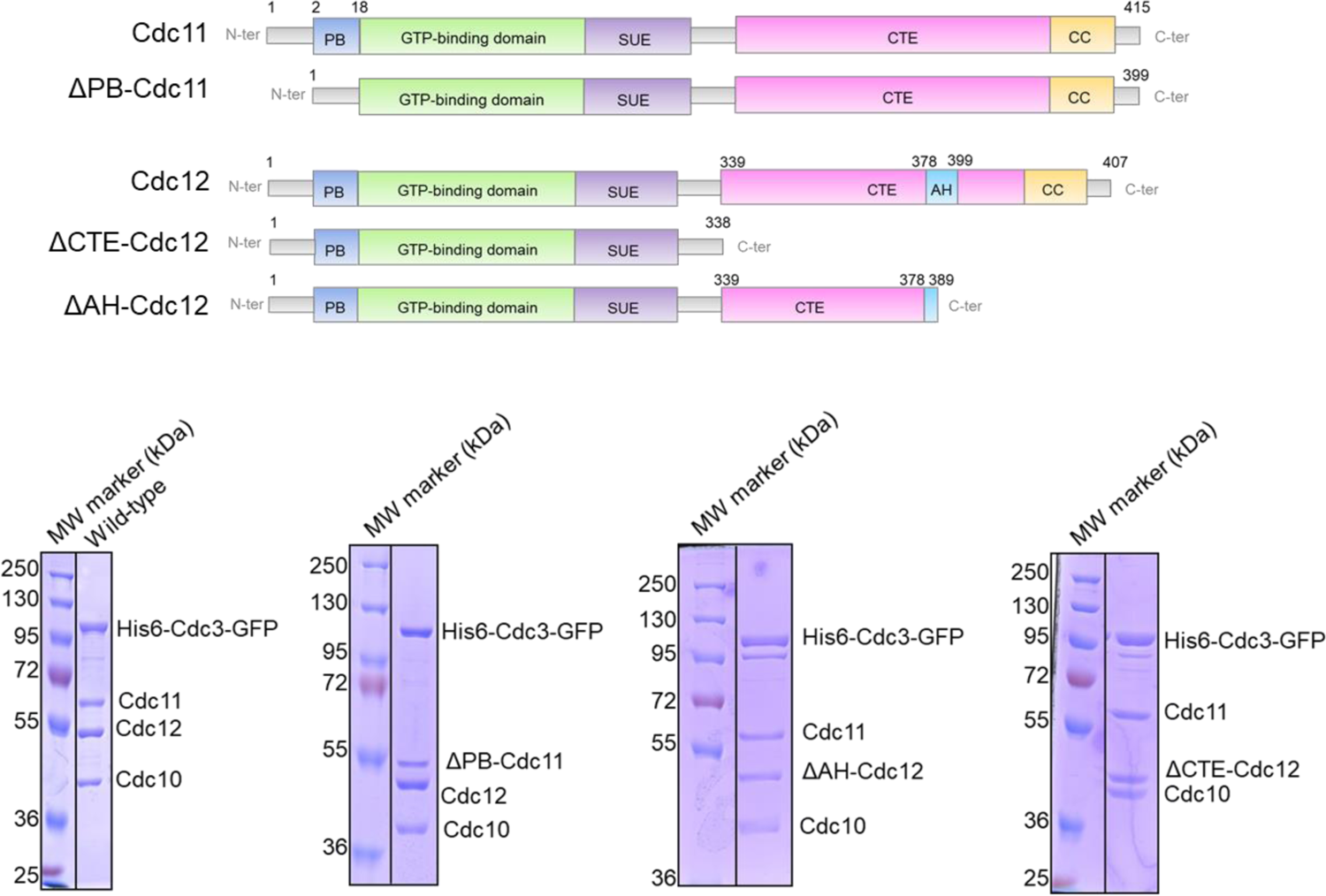
Domain representation of wild-type and mutant septins. Coomassie-stained SDS gel showing the purified GFP-tagged septin hetero-octamers used for *in vitro* assays.

**Figure 5-figure supplement 2.**
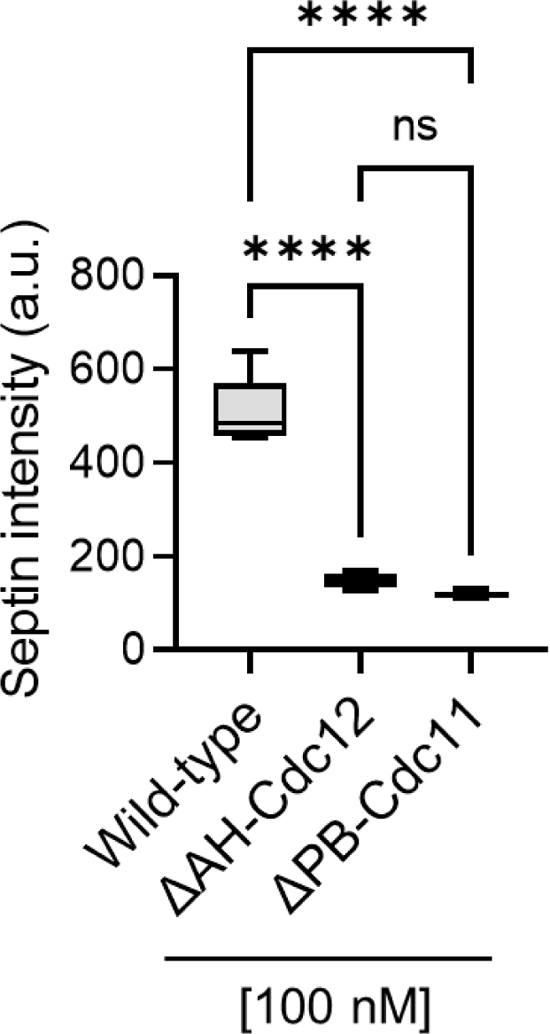
Mean gray value of 100 nM wild-type, ΔPB-Cdc11 or ΔAH-Cdc12 mutants septin-GFP complexes on the flat surface of silica nanostructured substrates functionalized with PI(4,5)P_2_-containing SLBs. Number of images analyzed, n, n = 56, 54 and 66 respectively, from two technical replicates. Error bars represent s.d..

**Figure 6-figure supplement 1.**
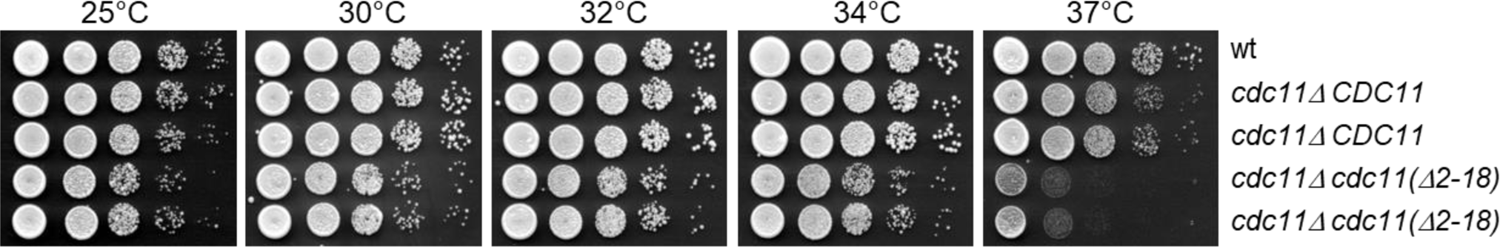
Viability of the *cdc11* (*Δ2-18)* mutant at different temperatures. Serial dilutions of cells with the indicated genotypes were spotted on YEPD plates and incubated at different temperatures (25°C, 30 °C, 32°C, 34°C, 37°C).

**Figure 6-figure supplement 2.**
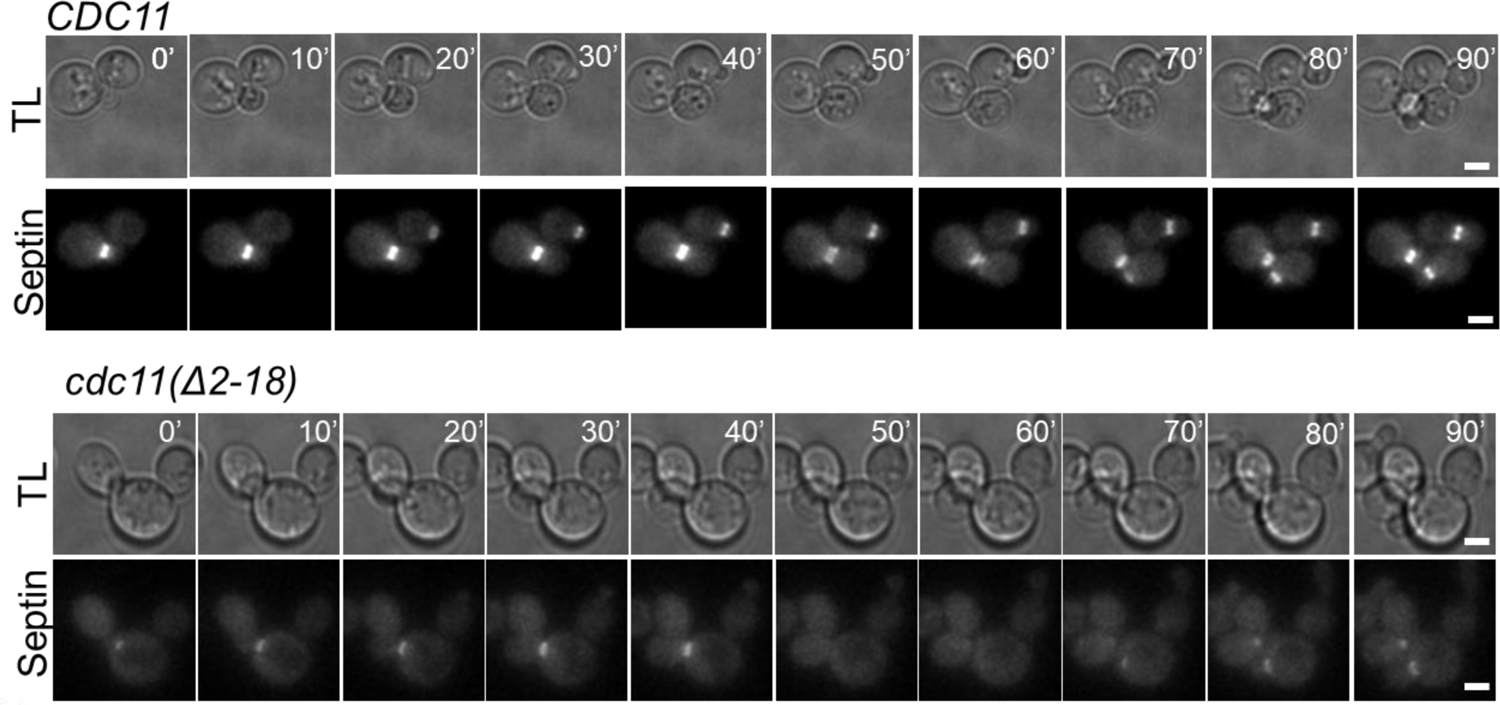
Transmitted light (TL) and TIRF images of wild-type *CDC11* and *cdc11(Δ2-18)* mutant cells expressing mCherry-Cdc3. Cells with the indicated genotypes were imaged every 5 min for 6 h at 37 °C. Scale bar, 2µm.

**Figure 6-figure supplement 3.**
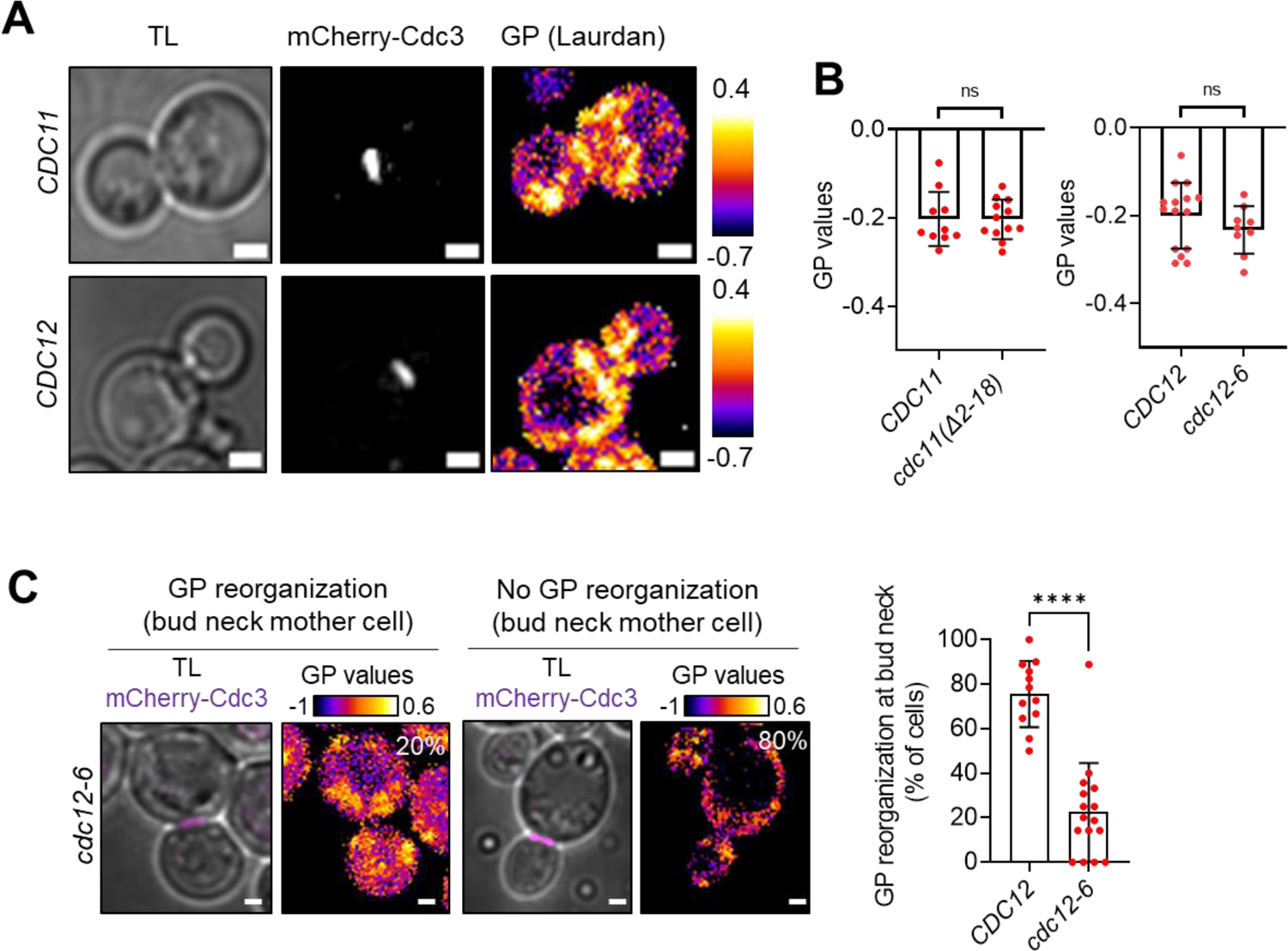
**A)** Transmitted light (TL) and confocal images of yeast cells expressing mCherry-Cdc3 in *CDC12* and *CDC11* cells stained with the Laurdan probe. Color-coded GP value images (fire LUT) of z-stacks max-projections. Scale bar, 2 µm. **B)** Quantification of the global GP index at the plasma membrane in *CDC12*, *cdc12-6, CDC11* and *cdc11(Δ2-18)* budded cells after staining with the Laurdan probe. Number of cells analyzed for each condition, n = 15, 7, 10, and 12, respectively, from three replicates. Error bars represent s.d.; Mann-Whitney test: n.s. P > 0.05. **C)** TL and confocal images of *cdc12-6* budded cells expressing mCherry-Cdc3 (magenta) and stained with the Laurdan probe. Color-coded GP value images (fire LUT) of z-stacks projections. Scale bar, 2 µm. % of *CDC12* and *cdc12-6* mutant cells displaying high GP values (GP > 1, indicative of Lo phases) at the bud neck of the mother cell cortex. Number of images analyzed, n = 12, and n = 16, respectively, from two independent experiments. Error bars represent s.d.; Mann-Whitney test: **** P < 0.0001.

